# Cell culture differentiation and proliferation conditions influence the *in vitro* regeneration of the human airway epithelium

**DOI:** 10.1101/2024.03.16.584842

**Authors:** Elisa Redman, Morgane Fierville, Amélie Cavard, Magali Plaisant, Marie-Jeanne Arguel, Sandra Ruiz Garcia, Eamon M McAndrew, Cédric Girard-Riboulleau, Kevin Lebrigand, Virginie Magnone, Gilles Ponzio, Delphine Gras, Pascal Chanez, Sophie Abelanet, Pascal Barbry, Brice Marcet, Laure-Emmanuelle Zaragosi

## Abstract

The human airway mucociliary epithelium can be recapitulated *in vitro* using primary cells cultured in an Air-Liquid Interface (ALI), a reliable surrogate to perform pathophysiological studies. As tremendous variations exist between media used for ALI-cultured human airway epithelial cells, our study aimed to evaluate the impact of several media (BEGM^TM^, PneumaCult^TM^, “Half&Half” and “Clancy”) on cell type distribution using single-cell RNA sequencing and imaging. Our work revealed the impact of these media on cell composition, gene expression profile, cell signaling and epithelial morphology. We found higher proportions of multiciliated cells in PneumaCult^TM^-ALI and Half&Half, stronger EGF signaling from basal cells in BEGM^TM^-ALI, differential expression of the SARS-CoV-2 entry factor *ACE2*, and distinct secretome transcripts depending on media used. We also established that proliferation in PneumaCult^TM^-Ex Plus favored secretory cell fate, showing the key influence of proliferation media on late differentiation epithelial characteristics. Altogether, our data offer a comprehensive repertoire for evaluating the effects of culture conditions on airway epithelial differentiation and will help to choose the most relevant medium according to the processes to be investigated such as cilia, mucus biology or viral infection. We detail useful parameters that should be explored to document airway epithelial cell fate and morphology.

## Introduction

The mammalian airways are lined by a mucociliary epithelium, composed of basal, club, goblet, multiciliated cells (MCCs), and rarer cell types (deuterosomal, ionocytes, pulmonary neuroendocrine and tuft cells, microfold cells) (1–3). The cell composition of the epithelium varies according to the macro-anatomical location, with the highest differences found between the nasal and the tracheobronchial epithelia (4). Within the tracheobronchial airways, cellular distribution is relatively stable, with modifications occurring in the most distal bronchioles (4–7). The airway epithelium is frequently challenged by inhalation of noxious compounds, chemicals, microorganisms and viruses, which alter its integrity. Specific repair mechanisms allow full epithelial restoration so that normal physiological function can be re-established (Supplementary Figure 1A). In chronic lung diseases, such as cystic fibrosis (CF), asthma or chronic obstructive pulmonary disease (COPD), frequent injuries and chronic inflammation induce a remodeling of the epithelium, often associated with a progressive loss of MCCs and an increased content of goblet cells. Studying the regeneration of the epithelium in normal or pathological conditions is necessary to identify mechanisms regulating the physiological regeneration and pathological remodeling of the airway epithelium. The development of cell cultures at the Air-Liquid Interface (ALI) of nasal or bronchial epithelial cells has been crucial in many mechanistic studies (8, 9). These cultures are not just convenient surrogates of *in vivo* epithelium (10–13): they were extensively used to establish the cellular roadmap of airway regeneration (10, 14) (Supplementary Figure 1A). This is illustrated by the recent approval of Kalydeco® by the Food and Drug Administration, based on work using primary cultures to predict clinical response in CF patients bearing ultra-rare CFTR mutations, for whom direct clinical trial studies are not feasible (15). Over the past three years, it also became the most appropriate model for SARS-CoV-2 *in vitro* infectivity studies. While most studies have identified MCCs as the major entry cell type (16–21), others have shown a more widespread distribution (22, 23), some with non-ciliated cells being infected preferentially over MCCs (24, 25). One striking difference between these studies was indeed the culture medium that was used.

Considering the large differences existing in cell culture conditions across the many studies using ALI cultures, it is important to define exactly the impact of the different experimental setups in term of cellular composition. Some studies have already performed direct comparison of cell culture media on ALI culture structure and/or function (26–33), but none has combined quantitative assessment of cell type distribution with differential gene expression profiles. The effect of using distinct commercial proliferation media during the cell expansion stage has never been reported either. We thus investigated the impact of 4 distinct proliferation and differentiation media on the epithelial structure and cellular composition of the reconstructed airway epithelium. We selected two commercially available and widely used media (PneumaCult^TM^ and BEGM^TM^) that we compared with “Half & Half”, which has been described by Susan Reynold’s group and which is considered as producing less biases compared to other media (34). The last medium is used by Clancy’s group to perform electrophysiological investigations (35). We used single-cell RNA sequencing (scRNA-seq) to quantify cell type distribution and gene expression profile variations across the 4 media. We also evaluated the impact of proliferation media on ALI differentiation, and the variations caused by the use of distinct porous membranes. Given the widespread use of ALI culture in SARS-CoV-2 studies, we also evaluated the effect of these media on viral entry factor expression.

## Methods

In order to evaluate the impact of cell culture media on cell composition of fully-differentiated ALI-regenerated airway epithelium, we first used primary cells that were freshly dissociated from human bronchi and set up proliferation and differentiation in 4 culture media including PneumaCult^TM^-ALI and BEGM^TM^-ALI, i.e. the two most widely used media, in parallel with Half & Half (34) and “Clancy” medium (35). Freshly isolated human bronchial epithelial (HBECs) cells underwent one passage in PneumaCult^TM^-Ex Plus medium, then were split and seeded on Transwell^TM^ membranes in 4 distinct proliferation media: PneumaCult^TM^-Ex Plus (hereafter named Pneuma-Ex+), BEGM^TM^ (BEGM), “Wu” medium and “Clancy” medium. Once cells reached confluence, differentiation was induced by removing the medium in the apical chamber and adding matched differentiation medium in the basal chamber. The differentiation media were PneumaCult^TM^-ALI (hereafter named Pneuma-ALI), Half & Half (H&H), BEGM^TM^-ALI (BEGM-ALI), and Clancy. Single-cell RNA sequencing (scRNA-seq) was performed at 3 time-points: after the initial propagation step on plastic flasks, at the onset of the air-liquid interface (ALI0) and at full differentiation (ALI28) (Supplementary Figure 1B). To evaluate whether some differences in ALI differentiation might stem from the medium used at the initial cell propagation steps, we amplified HNECs (Human Nasal Epithelial Cells) for 2 passages on plastic, in either BEGM or Pneuma-Ex+. We then seeded them on membranes, maintained cell amplification in the same medium until reaching confluence, and then set up the air liquid interface in either BEGM-ALI or Pneuma-ALI (Supplementary Figure 1C). We evaluated epithelial composition by quantitative PCR and immunostaining. The effects of using alternative semi-porous membranes were also evaluated. Detailed methods are described in the supplementary information file.

## Results

### ALI differentiation medium influences epithelial morphology and cell type distributions

After full differentiation of HBECs, epithelia were fixed for histological analysis and immunostainings. Hematoxylin/Eosin staining on epithelia sections revealed large differences in tissue structures (Figure 1A). Pneuma-ALI and H&H media generated thicker epithelia with an apical surface covered with cilia. BEGM-ALI and Clancy media generated much thinner epithelia, with this effect more prevalent for Clancy medium. Cilia immunostaining with acetylated alpha-tubulin suggested a higher content of multiciliated cells (MCCs) in Pneuma-ALI and H&H media than in BEGM-ALI and Clancy media (Figure 1A). MUC5AC+ goblet cell content was difficult to compare between culture media because staining patterns differed according to media with for instance smaller and more intense MUC5AC patches in Pneuma-ALI (Figure 1A).

**Figure 1.**
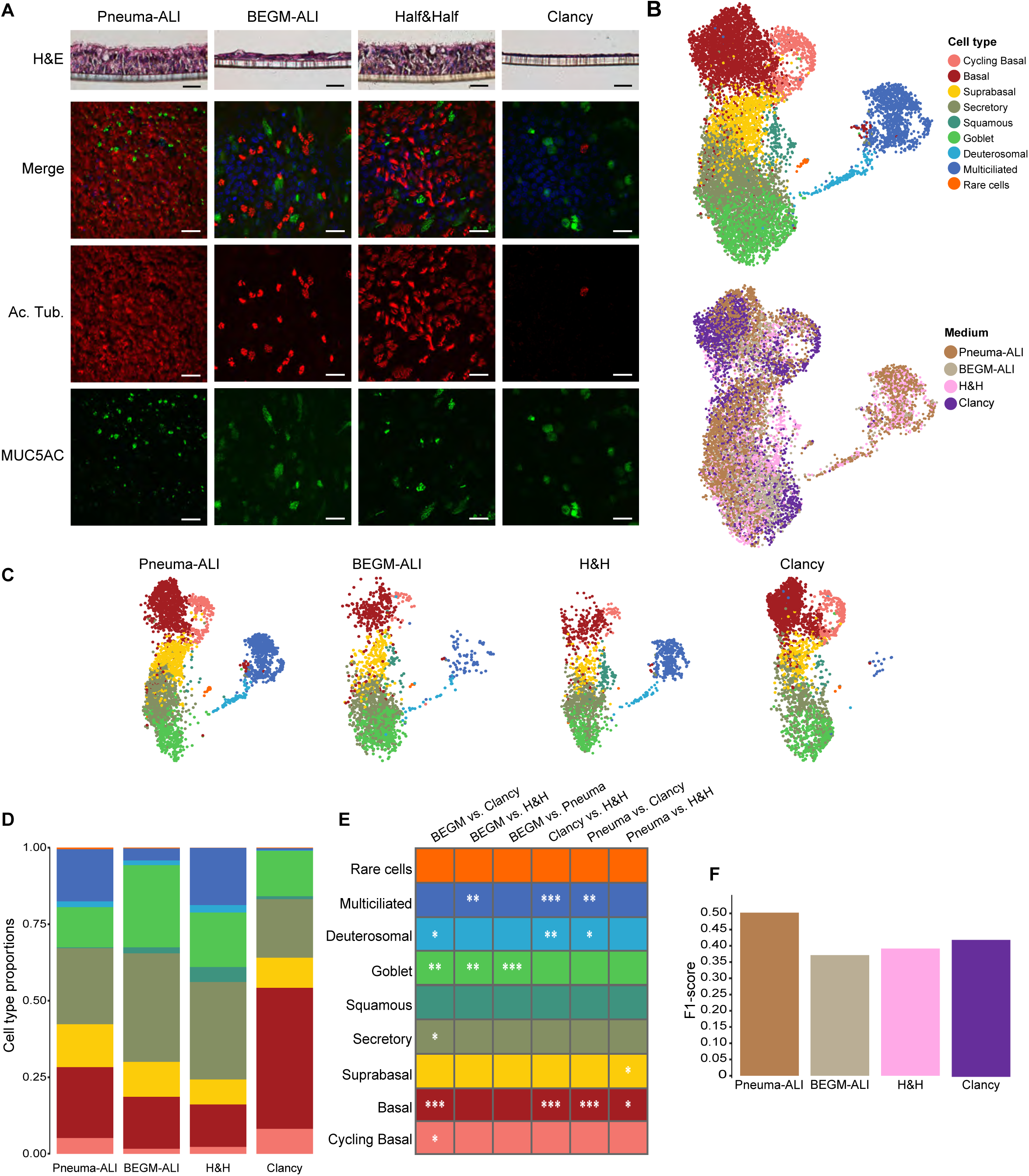
Single-cell RNA-seq analysis of human bronchial epithelial cells (HBECs) after 28 days of differentiation (ALI28) in 4 distinct media. **(A)** General characterization of HBEC differentiation in 4 cell culture media. Top panels: Representative images of Hematoxylin/Eosin staining of sections of HBECs at ALI28; Scale bars: 100 μm. For each condition and each of the 2 independent cell cultures, 2 inserts were sectioned and imaged. Bottom panels: Representative images of immunostaining of HBECs at ALI28 for acetylated alpha-tubulin and MUC5AC. Scale bars: 30 μm. Nuclei were stained with DAPI and shown in blue on the merged images. For each condition and each of the 2 independent cell cultures, 3 inserts were stained and imaged. **(B)** UMAP of the integrated dataset of HBECs at ALI28 containing all cells from the 4 distinct conditions, from 2 independent cultures from 2 healthy donors. Top panel: UMAP colored by cell type; bottom panel: UMAP colored by medium. **(C)** UMAPs for each medium, colored by cell type, with color code identical to (A). **(D)** Quantification of cell type proportions in each medium. **(E)** p-value of t-test by propeller package, using the asin transformation, for each 2-to-2 medium comparison of cell type proportions. **(F)** F1-score derived from Universal Cell Embeddings (UCE) label transfer against the Human Lung Cell Atlas (HLCA) core reference (1). The F1 score is calculated by comparing the predicted labels, obtained through the UCE label transfer method, against those manually annotated using marker genes and clustering. Maximal score is 1.

Thus, to obtain a more quantitative and detailed description of the regenerated epithelia in each culture medium, we performed 3’ scRNA-seq on fully differentiated HBEC cultures in each medium (ALI28). To reduce batch effects, we multiplexed all conditions using cell hashing (36). Datasets for all conditions and all donors were aggregated for subsequent analyses, allowing cell type quantification and differential gene expression between culture media (Figure 1B). Cell types were determined based on expression of known marker genes (Supplementary Figures 2A-C, Supplementary Table 1). Basal cells were identified by high expression levels of *KRT5* and *TP63* and cycling basal cells, by typical proliferation markers such as *MKI67*. Suprabasal cells were defined as cells located at the basal side of the epithelium, but not directly lying the basal lamina, expressing *KRT5* but very low *TP63*. Secretory cells included not only club cells that were *SCGB1A1*+ and *CYP2F1*+, but also other cell types that did not necessarily express *SCGB1A1*, but instead markers such as *AQP5* and *FAM3D*. Squamous cells were characterized by an intermediate signature between suprabasal and secretory cells, and also through expression of specific genes such as *SPRR1A*, *IVL* or *SCEL*. Goblet cells displayed a typical secretory cell gene expression program, with *SCGB1A1*, *BPIFA1*, *BPIFB1* along with the expression of *MUC5B* and/or *MUC5AC*. Of note, the length of the *MUC5AC* and *MUC5B* transcripts can affect their detection in scRNA-seq datasets, and probably contributed to an underestimation of *MUC5AC*+ and *MUC5B+* cells. MCCs were identified as *FOXJ1*+, and *DYNLRB2*+ while deuterosomal cells, which are cells amplifying centrioles, were *FOXJ1*+, *CDC20B+* and *DYNLRB2*-. Rare cells formed a small but distinct cell cluster, characterized by the expression of the ionocyte-specific *FOXI1*, as well as *HEPACAM2* and *NREP*, which we previously detected expressed by both ionocytes and neuroendocrine cells. We also detected *STMN1* a specific marker of tuft cells, and *MARCKSL1* and *CRYM* expressed by both tuft and neuroendocrine cells as expected (1, 4).

Figure 1B shows the distribution of the different cell type clusters at ALI28, after aggregation of the four datasets (top panel) and colored according to the differentiation media (bottom panel). The Uniform Manifold Approximation and Projection (UMAP) representation is split in Figure 1C according to the four media, with the corresponding cell type distributions quantified in Figure 1D. Pneuma-ALI and H&H cultures displayed similar distribution profiles (Figures 1C and 1D) with for instance 15.7% and 19.5% of MCCs respectively, as well as 13.1% and 17.7% of goblet cells, respectively (Figure 1D, Supplementary Table 2). BEGM-ALI and Clancy media cultures displayed more diverse profiles characterized by a much lower MCC content (3.8 % for BEGM-ALI and 0.6% for Clancy medium) and higher goblet cell proportion in BEGM-ALI (26.6%). We assessed the statistical significance of the differences between scRNA-seq conditions by performing pairwise comparisons with the Propeller package (37, 38) which confirmed the significance of the observations for the multiciliated and goblet cell populations (Figure 1E). The Clancy medium also yielded a significant difference in basal cell content against all other media (Figure 1E). When comparing to the luminal immunostainings from Figure 1A, the higher proportion of goblet cells and absence of significant decrease of MCCs in BEGM-ALI can be surprising, but can be explained by the greater tissue thickness in Pneuma-ALI and H&H. As the single-cell RNA-seq dataset quantifies all cells, from the basal to the luminal compartment, the proportion of luminal cells in thick epithelia, is decreased compared to thin epithelia. In addition, immunostainings detected MUC5AC+ cells only, as opposed to single-cell RNAseq which identified goblet cells based on the entire set of enriched genes, including *MUC5B*. We quantified goblet cells expressing *MUC5AC* alone, *MUC5B* alone or both and found some significant differences between differentiation media, with BEGM and H&H producing more *MUC5B*-only and *MUC5AC*-only goblet cells, respectively (Supplementary Figures 3A and 3B).

We next sought to compare the regenerated epithelia with *in vivo* data to identify the *in vitro* conditions that best reproduced the healthy airway environment. We used a label transfer tool based on Universal Cell Embeddings (UCE) (39) to map each medium-specific dataset onto the Human Lung Cell Atlas core reference (HLCA) (1). We computed an F1-score by comparing the predicted labels, obtained through the UCE label transfer method, against those manually annotated using marker genes and clustering. The F1-score evaluates the classification performance of the method and the method should perform best on cells most resembling the *in vivo* reference. Figure 1F shows that the best F1-score was obtained by Pneuma-ALI media (0.5), followed by Clancy (0.42), H&H (0.39) and BEGM (0.37). We used label transfer to obtain prediction proportions between our *in vitro* and the *in vivo* reference and we observed variable results depending on cell clusters (Supplementary Figure 4). Despite high mapping with HLCA basal cells, basal cells from BEGM-ALI and H&H also mapped with HLCA suprabasal cells, suggesting diverse gene expression profiles in these media. On the other hand, suprabasal cells from Pneuma-ALI and Clancy media better mapped onto their HLCA counterparts. MCCs from all media mapped almost perfectly with HLCA MCCs, but in Clancy and BEGM-ALI, MCCs also displayed high scores for deuterosomal cells, suggesting more immature MCCs in these media. Finally, the most striking difference was observed for Pneuma-ALI goblet cells which did not map with HLCA goblet cells. Instead, cells that we classified as secretory in Pneuma-ALI mapped with goblet cells from HLCA (Supplementary Figure 4). These data indicate that secretory and goblet cells, which share very close expression profiles (Supplementary Figure 2B) might carry subtle differences between media that influence their classification.

Thus, scRNA-seq allowed the identification of differences between differentiation media, with a higher MCC content in Pneuma-ALI and H&H, as well as gene expression differences shown by differential mapping to an *in vivo* reference. To investigate further these findings, we next aimed at identifying more specific effects of each differentiation medium on gene expression profiles within each cell cluster.

### ALI differentiation medium influences gene expression profiles

We first performed media pairwise comparison using a pseudobulk strategy. The largest number of differentially expressed genes were obtained when comparing Pneuma-ALI with BEGM-ALI and Clancy (Supplementary Figure 5 and Supplementary Table 3). In contrast, the lowest number of differentially expressed genes were observed when comparing H&H with Pneuma-ALI. We then displayed the top expressed genes in all pair-wise comparisons. Some genes were specifically enriched in one medium such as *IL33* and *ZBTB16* for Pneuma-ALI and Clancy, respectively (Figure 2A). *IL33* is expressed *in vivo* in healthy lung, in basal and suprabasal cells, as well as in endothelial cells (1, 4). Interestingly, it is a Th2-oriented cytokine that is involved in asthma susceptibility (40). *ZBTB16* encodes a zinc finger transcription factor that may play a role in the transcriptional memory of hormone stimulation (41). *CYP26A1* was also only expressed by basal cells from Pneuma-ALI medium. This cytochrome encoding gene contributes to retinoic acid clearing (42). As Pneuma-ALI is the only of the 4 media for which the composition is not publicly available, interpreting this difference concerning retinoic acid clearing was difficult. *VIM* was also enriched in cycling basal cells from Clancy medium, suggesting a mesenchymal-like phenotype. Pneuma-ALI and H&H shared several differentially enriched genes compared to the other media, such as the secreted peptides *TFF3*, *ELAPOR1*, *SCGB1A1*. They also shared the upregulation of *GLIPR2*, a Golgi-associated protein that negatively regulates autophagy (43). On the other hand, BEGM-ALI and Clancy media shared enriched genes such as several anion exchangers (*SLC5A5*, *SLC34A2*), secreted proteins (*STATH*, *VSTM2L*, *BPIFA2*) as well as *GCNT3*, an N-acetylglucosaminyltransferase contributing to mucin glycosylation. Altogether, these data suggest significant differences in the secretomes by Pneuma-ALI and H&H on one hand, and BEGM-ALI and Clancy on the other.

**Figure 2.**
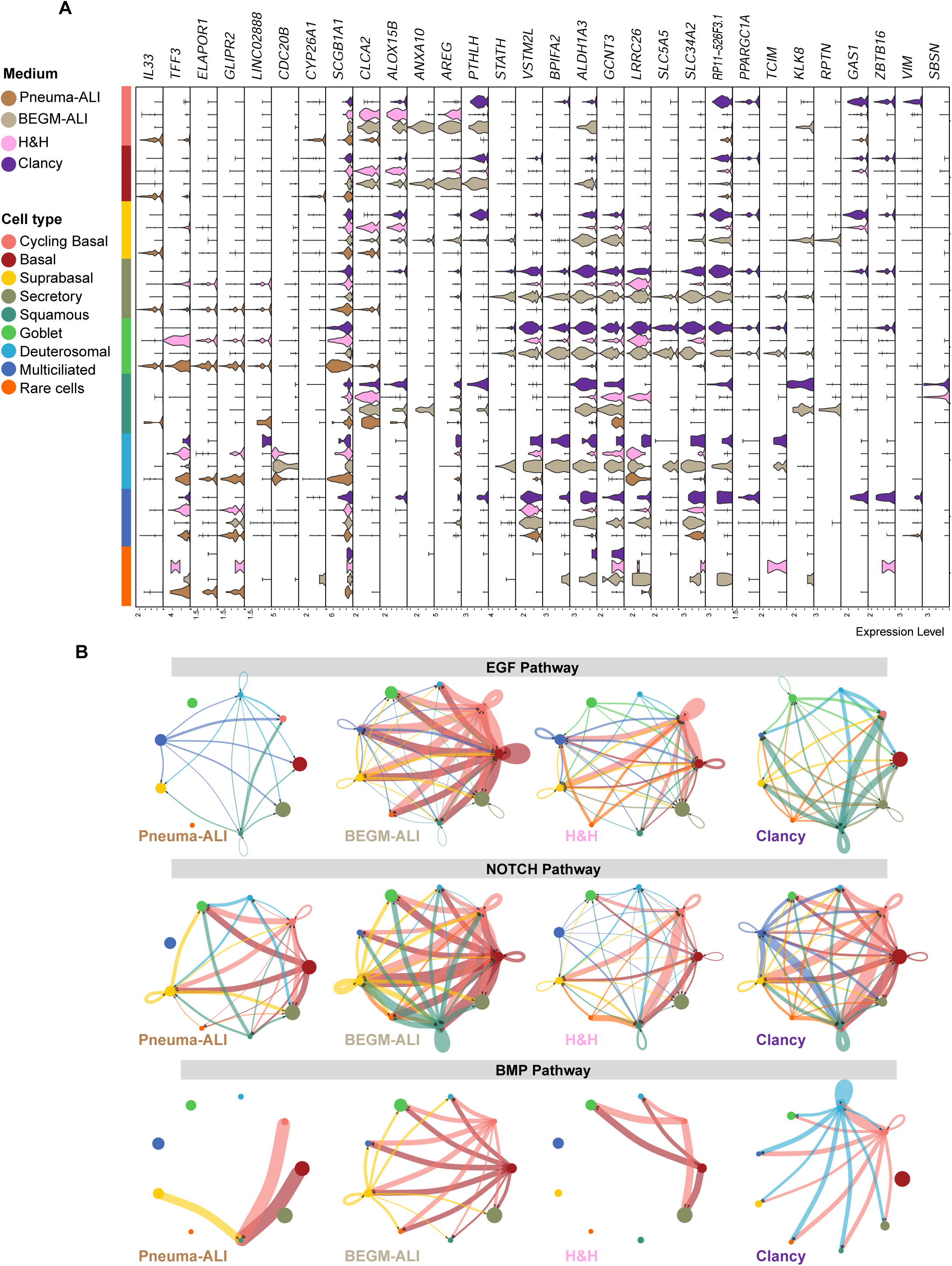
Genes and pathways regulated by each cell culture medium at ALI28 of bronchial epithelial cell differentiation. **(A)** Violin plot showing the top 30 regulated genes among all medium comparisons by pairs. Only genes expressed in at least 30% of cells of at least one cell cluster were selected. Top 30 genes were sorted by lowest adjusted p-value of each comparison by pair. For each gene, expression in the 4 tested media is shown. **(B)** Circle plots showing the inferred intercellular communication networks for EGF, NOTCH and BMP signaling. The edge width is proportional to inferred interaction strengths considering ligand-receptor pairs. Black arrowheads show directionality of the inferred interactions. For (B), the color code for cell types is identical to (A).

Given the tremendous number of possible comparisons, we used CellChat to stratify the different expressed pathways. CellChat quantifies outgoing and incoming signals per signaling pathway, at the level of each cell cluster, thus identifies cells involved in sending and receiving signals for each pathway, in an autocrine and paracrine manner. When comparing the number of inferred interactions within each culture medium, we found that BEGM-ALI generated the highest number of interactions compared to all other media, and that basal and suprabasal cells are the source of the major incoming signals (shown by top bars of incoming signaling patterns on Supplementary Figure 6, and by Supplementary Figure 7A and 7B). However, the communication probability of these interactions, represented by the interaction strength or "weight” calculated by CellChat, appeared equivalent between all media (Supplementary Figure 7A right panel and 7B). Some pathways showed striking differences in communication probabilities, such as the EGF, NOTCH, BMP and WNT pathways (Figure 2B and Supplementary Figure 7C). The EGF pathway showed stronger outgoing signals in BEGM-ALI and H&H media (Supplementary Figure 6) with basal and cycling basal cells sending EGF signals to all other cell types of the epithelium, including on themselves (Figure 2B). This signal is consistent with the strong expression of *AREG* in cycling basal and basal cells of BEGM-ALI and H&H, together with the specific *EREG* and *HBEGF* signals in BEGM-ALI only (Supplementary Figure 8A). Contribution of each Ligand-Receptor pair could be inferred from the expression data and showed high contribution of *AREG* signaling towards *EGFR* or *EGFR* together with *ERBB2* in BEGM-ALI (Supplementary Figures 8B-C). We also looked more closely at differences observed for the NOTCH pathway. Expression of ligands and receptors were modulated by differentiation media (Supplementary Figure 9A) which produced large differences in the inferred interactions (Supplementary Figures 9B-C). We found that suprabasal, deuterosomal, multiciliated and squamous cells were predicted as receiving high Notch pathway signals in BEGM compared to Pneuma-ALI (Supplementary Figure 9C). Since AREG has been reported to increase cell proliferation and mediate upregulation of mucus-related genes in the mouse lung and airway epithelial cell lines (44–47) and that the NOTCH pathway favors the secretory fate over the multiciliated fate (48–51), the differences we report here might explain the higher number of goblet cells and lower number of MCCs detected in BEGM-ALI.

Hence, the choice of differentiation media can strongly influence signaling pathways between the distinct cell types of the epithelium, which might affect the balance between the different cell types.

### Identification of cell types at the onset of differentiation

As differentiation media influenced epithelial composition and gene expression, we next assessed whether the cell composition at the onset of differentiation was equivalent between all culture conditions.

We first analyzed HBECs from the same donors as previously, immediately after the initial amplification step in PneumaCult^TM^-Ex Plus (Pneuma-Ex+) (Supplementary Figure 1B) to identify the different cell identities that were seeded on the culture membranes. Among the 1716 cells we analyzed, we found a majority of basal and cycling basal cells (Supplementary Figure 10, Supplementary Table 4). Interestingly, *KRT13*+ basal cell types were identified and comprised some proliferative cells, based on *MKI67* expression. A small fraction (3.50%) of goblet cells was also detected, identified by the expression of *SCGB1A1* and *MUC5AC*. No proliferation was detected among goblet cells, as evidenced by the absence of *MKI67* expression (Supplementary Figures 10A and 10B).

We then evaluated whether the use of the 4 distinct proliferation media during the HBEC propagation step on Transwells^TM^ membranes could influence epithelial cell composition at the starting point of air-liquid interface (ALI0), using scRNA-seq. Although basal and cycling basal cells composed the majority of the cultures, we also detected secretory cells (*SCGB1A1*+, *SLPI*+), and suprabasal cells with a secretory signature (*LY6D*+, *SLPI*+) (Figures 3A and 3B). No *MUC5AC*+ goblet cells were detected at this stage. Few cells that we named “undefined” were detected predominantly in the BEGM condition and did not match with any cell type usually found *in vivo.* These cells are negative for all basal cell markers, and positive for *DDIT3*, *SLC3A2*, *ISG15*, *SQSTM1*, among other specifically expressed genes (Figures 3A-C, Supplementary Table 5) that seem related to DNA damage, ER stress, negative regulation of RNA transcription and protein ubiquitination. Cell composition was affected by proliferation medium: the BEGM medium generated a smaller fraction of proliferative cells, which is consistent with the lower cell densities observed in this medium, in spite of reaching confluence. Indeed, contact inhibition appeared higher in BEGM (data not shown). BEGM generated more secretory cells, with lower level of expression of *SCGB1A1* than all other media (Figures 3B and 3C). In the absence of proliferation in these secretory cells, they probably emerged from differentiation during the expansion process. Wu and Pneuma-Ex+ generated similar cell compositions (Figure 3C). We noticed at this stage a ubiquitous expression of the goblet cell-specific transcription factor *SPDEF* in Pneuma-Ex+ and Wu medium (Figure 3B). *SPDEF* was not detected in either BEGM or Clancy medium in basal, cycling basal or suprabasal/secretory intermediates, but was detected in secretory cells from Clancy medium. To explore further this finding, we set up cultures on 4 independent batches of human nasal epithelial primary cells (HNECs), from 4 distinct healthy donors, either in BEGM or in Pneuma-Ex+. HNECs were used at this stage as a convenient material for experimental replication and were shown as reliable surrogates to HBECs (9). Quantitative PCR analysis confirmed strong upregulation of *SPDEF* in all 4 HNEC cultures at the proliferation stage (Supplementary Figure 10C). We then repeated this experiment with 3 additional HNEC cultures either in Pneuma-Ex+ or PneumaCult^TM^-Ex (Pneuma-Ex), which is equivalent to BEGM for proliferation and performed scRNA-seq analysis. In this dataset, we could identify cell populations that were similar to those we found in HBECs, and detected few multiciliated and endothelial cells, which were probably carried over from tissue isolation (Figure 3D, Supplementary Figure 10D, Supplementary Table 6). We confirmed that proliferation in Pneuma-Ex+ for one passage was sufficient to induce *SPDEF* expression (Figure 3E). Surprisingly, we found few proliferative goblet cells at this stage of cell propagation, with a clear enrichment in Pneuma-Ex+ compared to Pneuma-Ex (Figure 3F).

**Figure 3.**
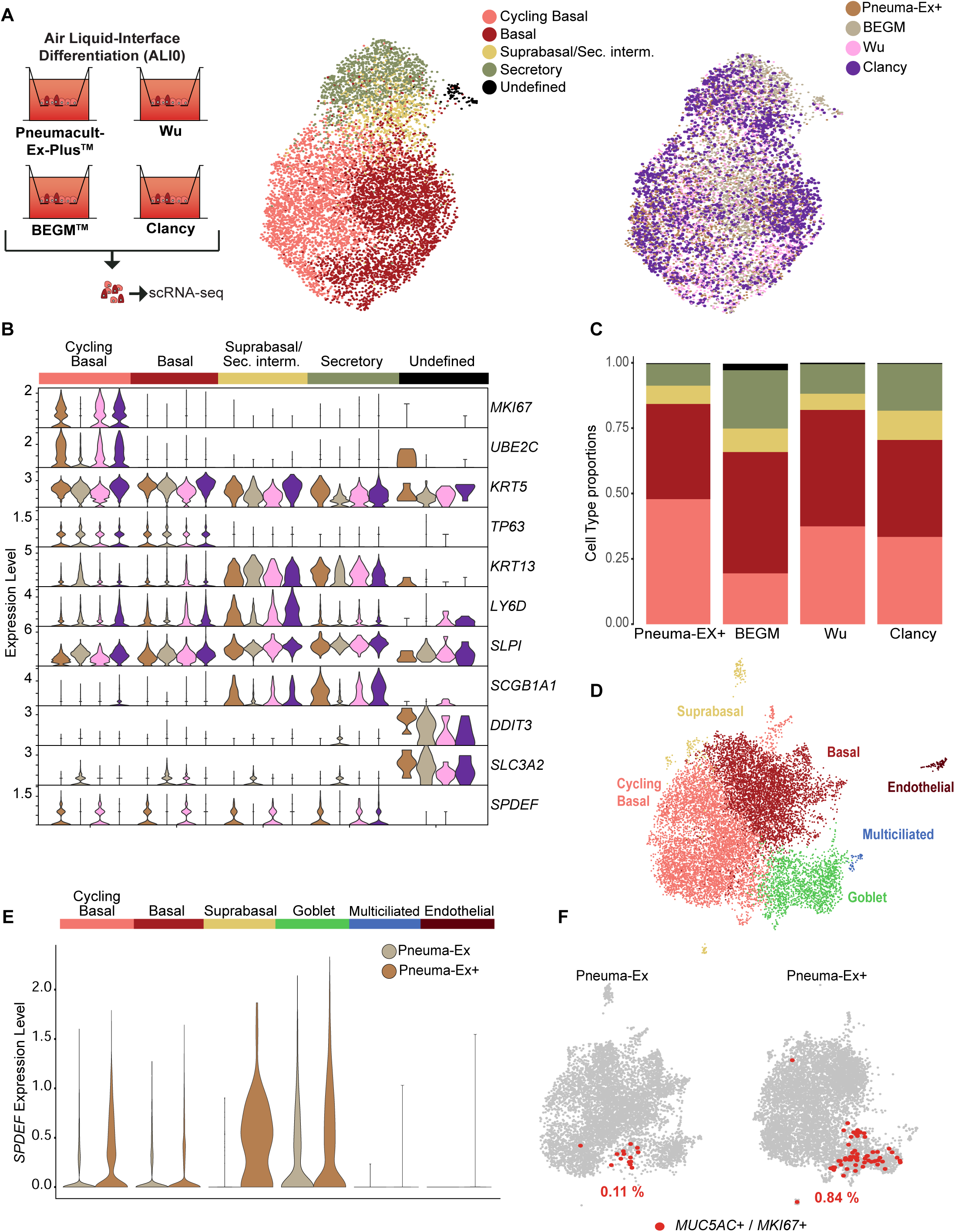
Impact of distinct proliferation media on cell type composition and cell expression at ALI0 of epithelial cell differentiation. **(A)** Left panel: experimental strategy. Right panel: UMAPs of the integrated dataset at the onset of differentiation of HBECs containing all cells from the 4 distinct conditions, colored by cell type and by medium according to the indicated color codes. **(B)** Expression of the top 10 marker genes and of *SPDEF*, for the 5 detected cell populations from (A), displayed for each proliferation medium. **(C)** Quantification of cell type proportions from (A) for each proliferation medium. **(D)** UMAP of the integrated dataset composed of HNECs analyzed after the first expansion stage, in either Pneuma-Ex or Pneuma-Ex Plus, colored by cell type. **(E)** Expression of *SPDEF* by each cell type of dataset from (D), in either Pneuma-Ex or Pneuma-Ex+ as indicated by color code displayed in (D). **(F)** Identification of *MUC5AC*+/*MKI67*+ cells in the integrated dataset composed of HNECs analyzed after the first expansion stage, in either Pneuma-Ex or Pneuma-Ex Plus. Cells were selected if they had normalized expression of *MUC5AC*>1 and *MKI67*>0.5. The percentage of *MUC5AC*+/*MKI67*+ cells for each sample is indicated on the plot.

Altogether, these results indicate that proliferation conditions could affect cell type distribution and gene expression profiles. Thus, we next evaluated whether some differences in ALI differentiation might stem from the initial cell propagation step.

### Effect of proliferation medium on features of differentiated cultures

HNECs were amplified in either BEGM or Pneuma-Ex+, on plastic flasks and on Transwells^TM^ membranes, and then induced to differentiate at the air-liquid interface in either BEGM-ALI or Pneuma-ALI (Supplementary Figure 1C). We evaluated epithelial composition by quantitative PCR and immunostaining using the MCC specific markers *FOXJ1* and acetylated alpha-tubulin, and the goblet cell specific marker *MUC5AC*. Both qPCR and immunostainings showed that although Pneuma-ALI yielded more MCCs than BEGM as expected, the proliferation medium had no effect on the MCC content. However, when differentiation was performed in BEGM-ALI, the use of Pneuma-Ex+ for proliferation increased MUC5AC content as shown by qPCR (Figure 4A), immunostainings (Figure 4B) as well as Western blot (Figures 4C and 4D, Supplementary Figure 11A-B). When differentiation was performed in Pneuma-ALI, although qPCR did not show any difference in MUC5AC expression comparing proliferation in BEGM or Pneuma-Ex+ (Figure 4A), Western blot did show a significant increase in MUC5AC content when cells were amplified in Pneuma-Ex+ (Figures 4C and 4D, Supplementary Figures 11A-B). MUC5B content tended to also be increased by proliferation in Pneuma-Ex+, although not significantly (Supplementary Figures 11C-E).

**Figure 4.**
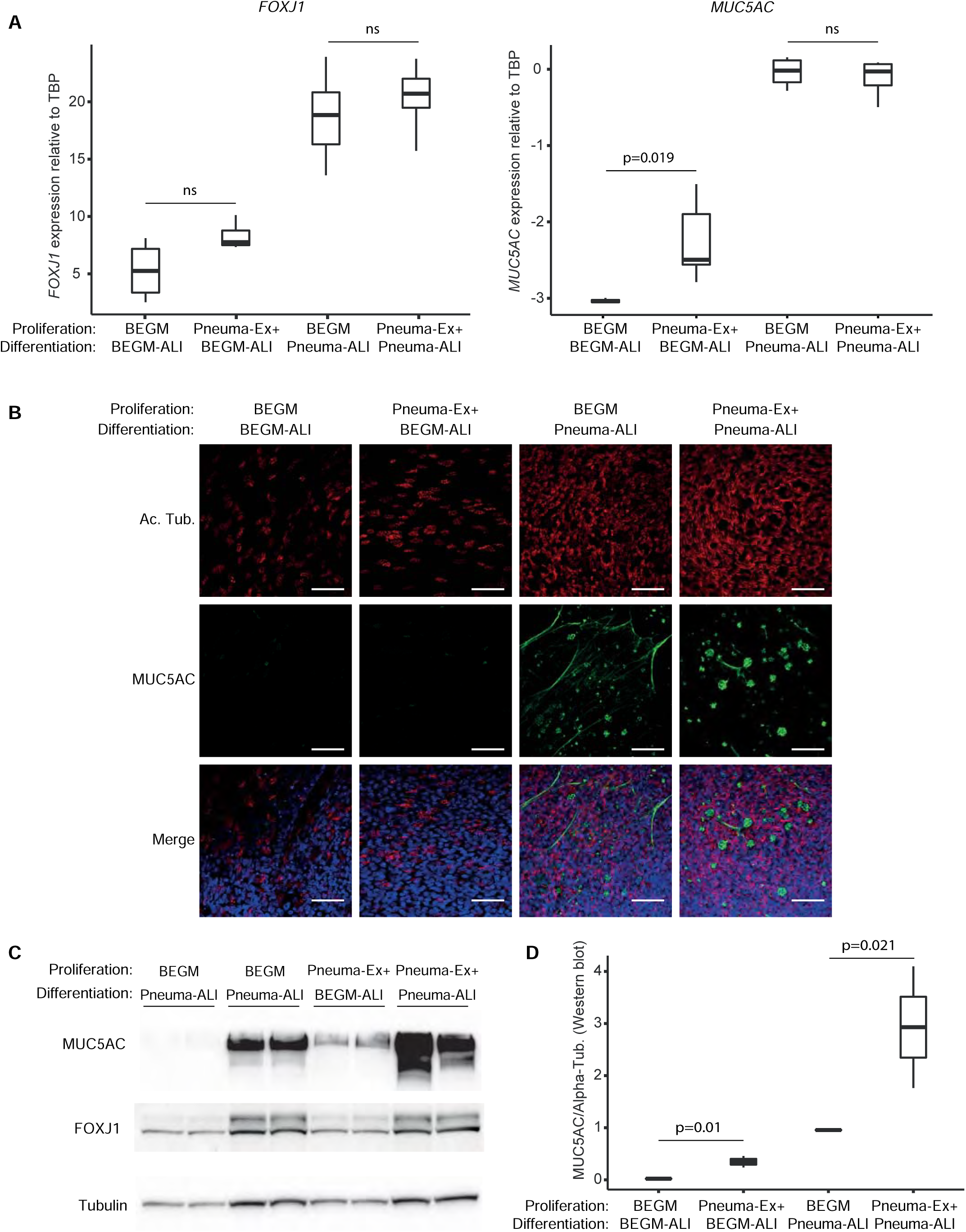
Effect of distinct proliferation media on epithelial composition after ALI differentiation of nasal epithelial cultures. **(A)** Expression of *FOXJ1* and *MUC5AC* (qPCR) on ALI cultures after full differentiation of nasal epithelial cells in the indicated media. Data shown are the values for 2^Ct(*FOXJ1*-*TBP*) and log10(2^Ct(*MUC5AC*-*TBP*)) for 3 independent cultures from 3 donors. p-values are the results of Mann-Whitney U test. **(B)** Representative images for immunostaining of MUC5AC and acetylated alpha-tubulin on HNEC ALI cultures after full differentiation in the indicated media; scale bars: 30 µm. Nuclei were stained with DAPI and shown in blue on the merge images. For each condition, 3 independent cell cultures were performed, and 3 inserts were stained and imaged for each. **(C)** Western blot for MUC5AC and FOXJ1 on HNEC ALI cultures after full differentiation in the indicated media. Tubulin is used as loading control. For each condition, two lanes were loaded, each with an independent Transwell^TM^ membrane from the same culture. **(D)** Quantification of the MUC5AC signal from (C), p-values are the results of Welch’s t-test. For each condition, 3 independent cell cultures were performed, and 2 inserts were used for Western Blot. Data from the additional cultures are shown on Supplementary Figure 11.

Thus, even though proliferation in Pneuma-Ex+ did not favor a goblet or secretory cell content during proliferation (Figure 3, Supplementary Figure 10D), this proliferation medium favored goblet cell differentiation or mucin production. This increase might be due to the proliferative goblet cells that we detected during the initial cell propagation of nasal cultures in Pneuma-Ex+ (Figure 3F) although there is no evidence that these cells were retained during the amplification step on Transwell^TM^ membranes. For HBECs, even though we detected some goblet cells at the initial propagation step (Supplementary Figures 10A and 10B), these cells were not retained after amplification on membranes (Figure 3A). Another possibility is that Pneuma-Ex+ induced basal cells imprinting towards a subsequent goblet fate as suggested by increased expression of SPDEF (Figures 3B and 3E).

### Effect of cell culture media on SARS-CoV-2 entry factors

The ALI model is appropriate for SARS-CoV-2 *in vitro* infectivity studies but discrepancies in major entry cell types have been described (16–24). As various cell culture media have been used in past studies, we wondered whether SARS-CoV-2 entry factor expression varied according to differentiation and proliferation media. While *ACE2* was very poorly detected in scRNA-seq data of HBECs at ALI28 (Figure 5A), other entry factors such as *TMPRSS2*, *FURIN*, *BSG*, *CTSB*, *ANPEP*, *ST6GAL1* and *ST3GAL4* were well detected.

**Figure 5.**
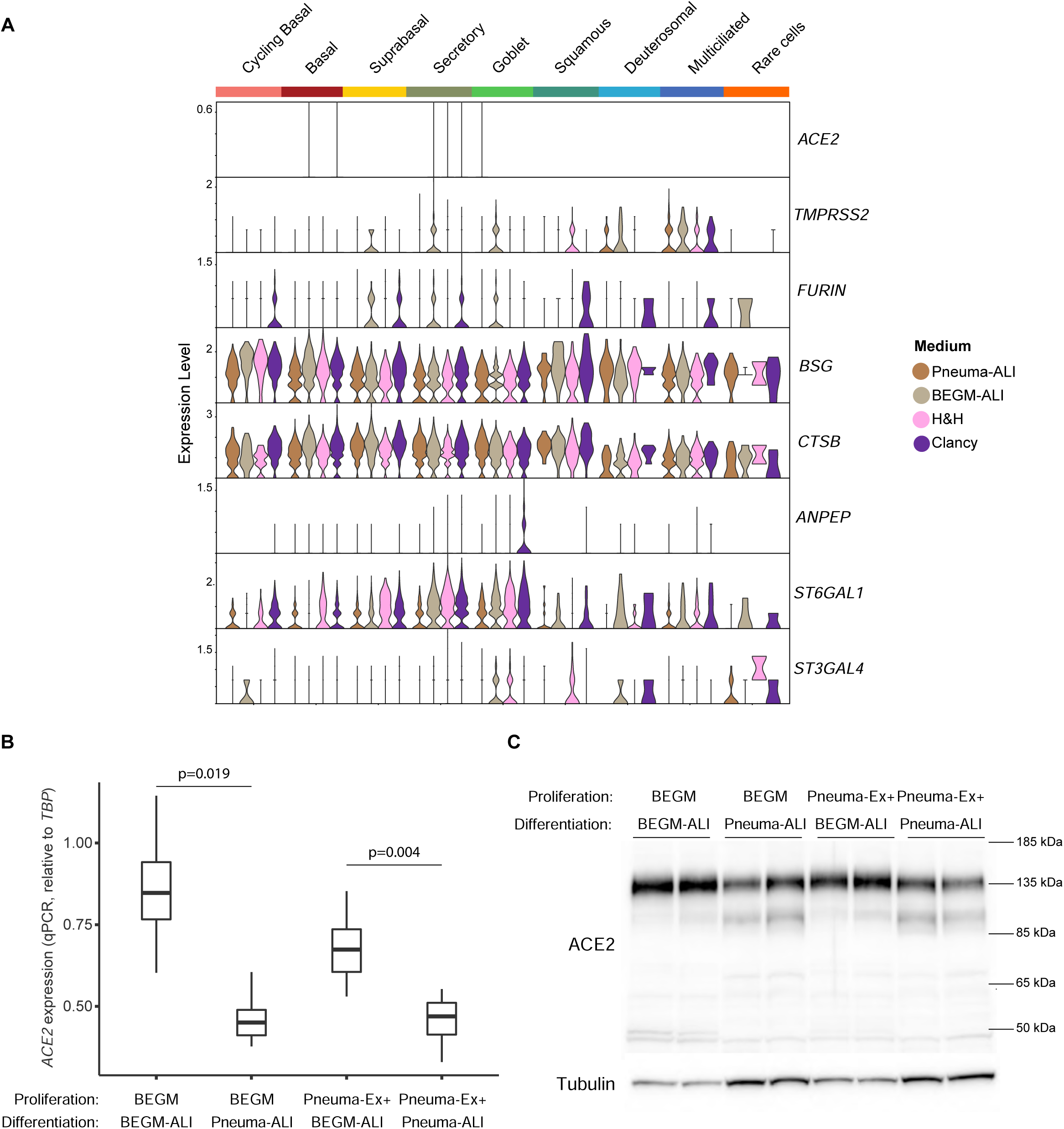
Effect of proliferation and differentiation media on ACE2 expression at ALI28 of epithelial cell differentiation. **(A)** Violin plot showing expression of SARS-CoV-2 entry factors in each of the 4 media conditions used for HBECs, at ALI28 (Figure 1B). **(B)** qPCR for *ACE2* on 3 independent HNEC ALI cultures from 3 donors after proliferation and full differentiation in the indicated media. Data shown are the values for 2^Ct(*ACE2*-*TBP*) for 3 independent cultures from 3 donors. p-values are the results of Mann-Whitney U test. **(C)** Western blot for ACE2 on HNEC ALI cultures after full differentiation in the indicated media. Tubulin is used as loading control. For each condition, 3 independent cell cultures were performed, and 2 inserts were used for Western Blot. Data from 2 additional independent cultures are shown on Supplementary Figure 12.

*TMPRSS2* was well expressed in MCCs from all media but was only detected in BEGM-ALI for secretory and goblet cells and in H&H for squamous cells. Other entry factors showed some media-specific expression such as *FURIN* in BEGM-ALI and Clancy medium, *ANPEP* which was restricted to Clancy medium in goblet cells and *ST3GAL4* which was detected in BEGM-ALI and H&H for goblet cells, and H&H only in squamous cells (Figure 5A). *ACE2* expression was then investigated by qPCR and western blot, for the two proliferation and differentiation commercial media, in HNECs. Although proliferation media did not produce significant effects (Figures 5B and 5C, Supplementary Figure 12A), qPCR showed that ACE2 expression was significantly lower in Pneuma-ALI. The Western blot analysis showed a decreased expression of the main isoform in Pneuma-ALI compared to BEGM-ALI which appears as a ∼135 kDa band, and an increased expression of a ∼100 kDa differentially glycosylated isoform (52) (Figure 5C, Supplementary Figure 12B).

Thus, both ACE2 expression and glycosylation displayed media specific effects.

### Effect of membranes and media alternatives

The pandemic crisis and the development of ALI models in respiratory virus studies led to reagent shortages. Alternative media or porous membranes other than Transwell^TM^ were evaluated. PneumaCult^TM^-ALI-S (Pneuma-ALI-S), an additional commercial medium, designed for the differentiation of small airways, and Thincerts^TM^ membranes (Greiner Bio-One), a commercial membrane, were assessed on HNECs. Morphology of multiciliated and goblet cell and content in reconstructed epithelium were compared between Pneuma-ALI-S and Pneuma-ALI. Figure 6 and Supplementary Figure 13 show that the use of the Pneuma-Ex+/Pneuma-ALI combination appeared to yield more MUC5AC+ cells than the Pneuma-Ex/Pneuma-ALI combination. In addition, when comparing epithelial layer morphology, the use of Pneuma-Ex+ during the proliferation phase, generated thick epithelia, with invaginations within the layer, as shown by actin, hematoxylin-eosin as well as nuclei staining (Figure 6 and Supplementary Figure 13), which is not the case when cells proliferate in Pneuma-Ex. In contrary, in spite of the use of Pneuma-Ex+ for proliferation, the use of Pneuma-ALI-S restored the epithelial morphology, as shown by the absence of invaginations in the reconstructed tissue, and a flat epithelial surface. MUC5AC+ cell content appeared similar to that obtained with Pneuma-ALI (Figure 6 and Supplementary Figure 13). Even though further experiments are needed to precisely evaluate the effects of the use of Pneuma-ALI-S on cell composition of ALI cultures, this medium can be used to generate apparent healthy epithelia after HNEC expansion in Pneuma-Ex+. We also evaluated the effect of using alternative polyethylene terephthalate (PET) membranes which are very similar to Transwell^TM^ filters that carry 0.4 µm pores. The difference between these products stands in the pore density which is 2.10⁶ pores per cm² for Thincerts^TM^ and 4.10⁶ pores per cm² for Transwells^TM^. We evaluated these membranes after HNEC proliferation in Pneuma-Ex+ medium, as above, and induced differentiation with either Pneuma-ALI or Promocell®-ALI (Promo-ALI, which composition is identical to BEGM-ALI). As previously mentioned, thick and invaginated epithelia were obtained when using Transwell^TM^ membranes with the Pneuma-Ex+/Pneuma-ALI combination (Figure 7). Promo-ALI generated thinner epithelia, but they still displayed invaginations. The basal and suprabasal cell layers, stained with KRT5 seemed expanded in both media. The use of Thincert^TM^ membranes strongly reduced the overall thickness of the epithelia and yielded epithelia with the expected morphology together with the expected basal, goblet and MCC disposition in Pneuma-ALI. Nonetheless, the use of Promo-ALI on Thincert^TM^ membranes produced thin epithelia containing few multiciliated and MUC5AC+ cells (Figure 7).

**Figure 6.**
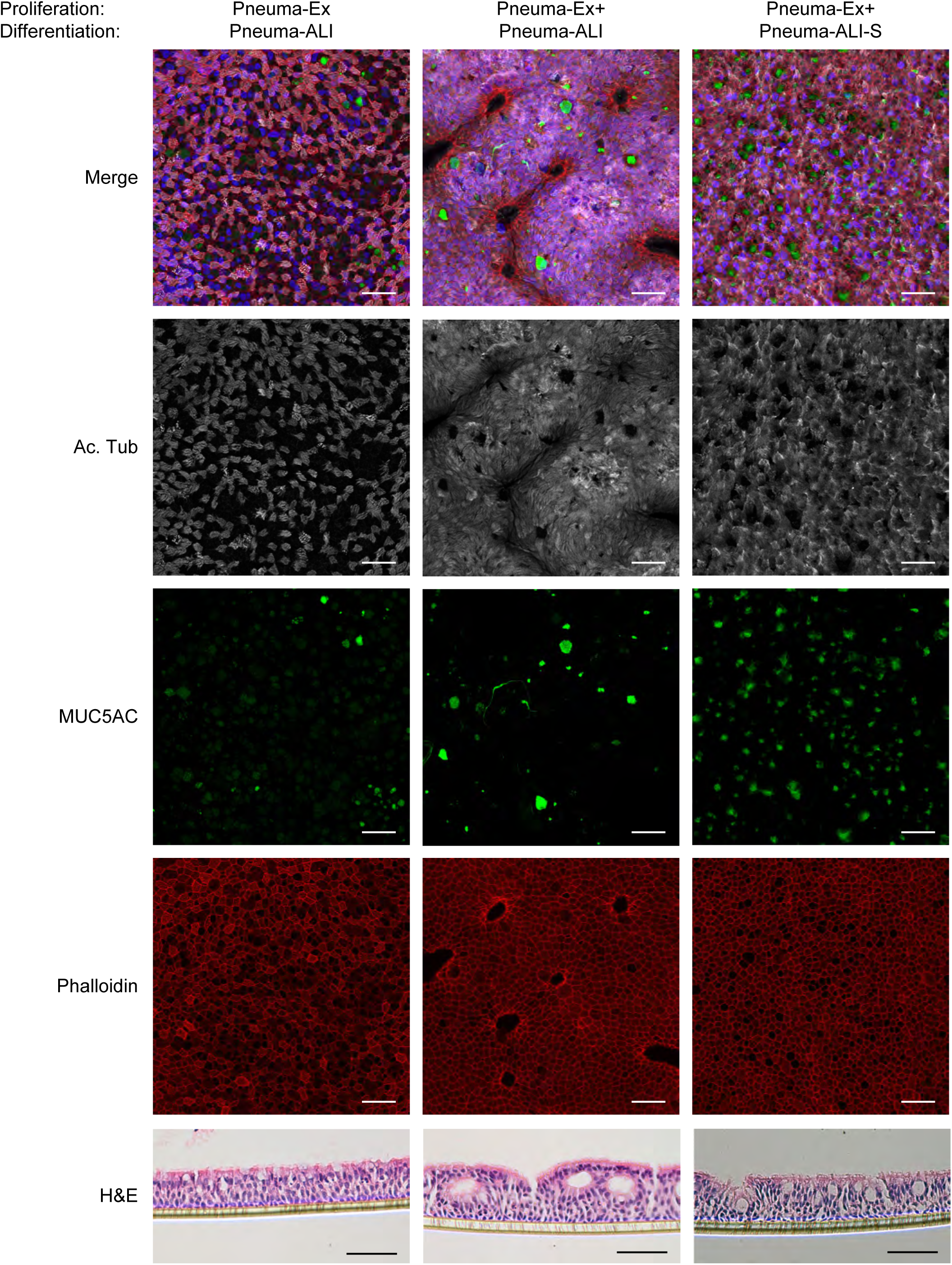
Effect of an alternative differentiation medium on epithelial composition after ALI differentiation of nasal epithelial cells. **Top panels:** Representative images of immunostaining for MUC5AC and acetylated alpha-tubulin on HNEC ALI cultures after full differentiation in the indicated media. Phalloidin was used to stain for actin. Nuclei were stained with DAPI and shown in blue on the merge images; scale bars: 30 µm. **Bottom panels:** Hematoxylin/Eosin staining of sections of paraffin-embedded epithelia; scale bars: 100 µm. For each condition, one cell culture was performed, and 2 inserts were stained and imaged. Additional images are shown in Supplementary Figure 13.

**Figure 7.**
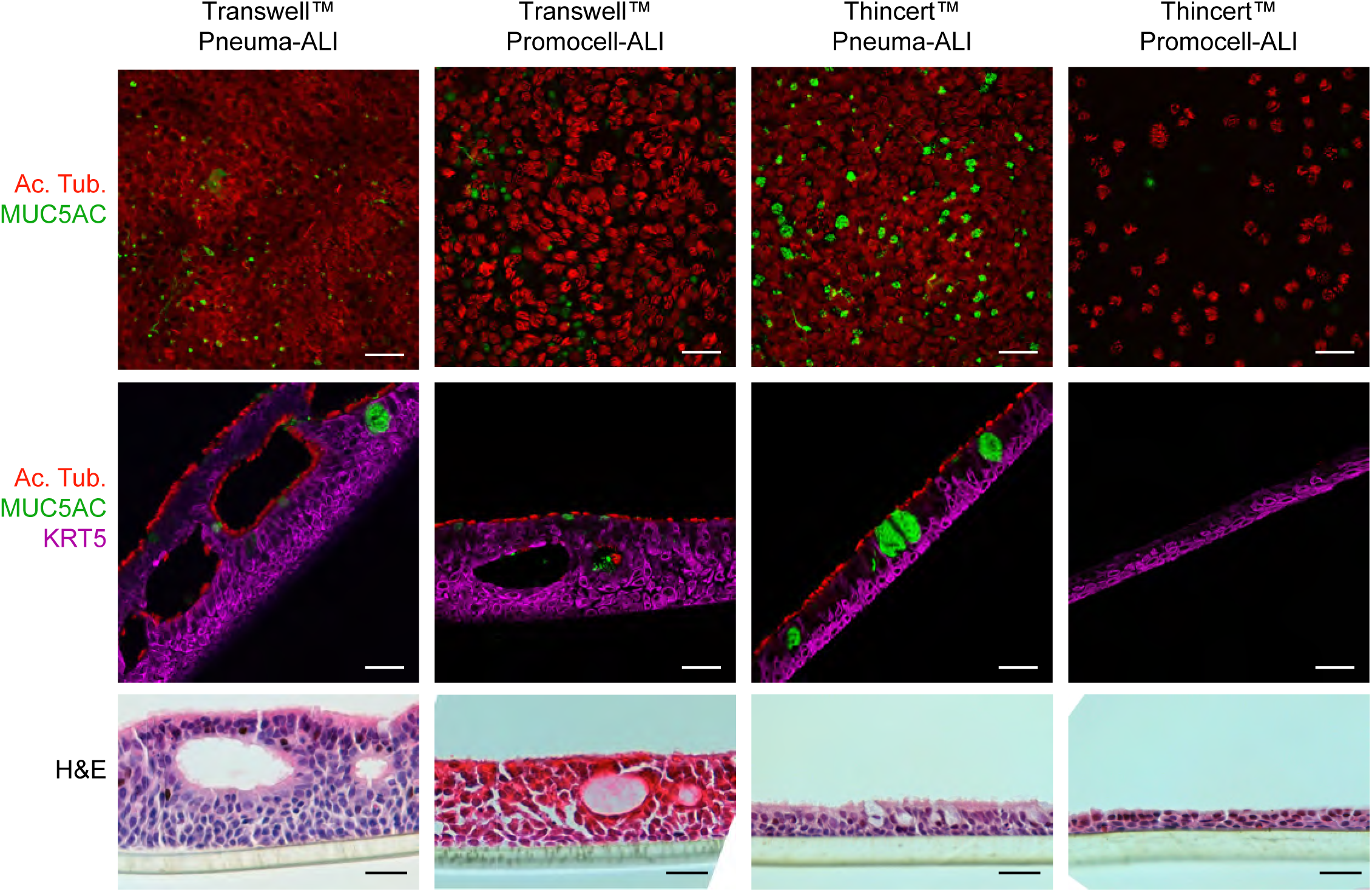
Effect of cell culture inserts and differentiation media on epithelial composition after ALI differentiation of nasal epithelial cultures. **Top panels**: Representative images of immunostaining on whole inserts for MUC5AC and acetylated alpha-tubulin on HNEC ALI cultures after full differentiation in the indicated media; scale bars: 30 µm. **Middle panels:** Representative images of immunostaining on sections of inserts for MUC5AC and acetylated alpha-tubulin on ALI cultures after full differentiation in the indicated media; scale bars: 30 µm. **Bottom panels:** Representative images of Hematoxylin/Eosin staining of sections of paraffin-embedded inserts; scale bars: 100 µm. For each condition, 2 inserts were stained and imaged.

Together, these results show that if using Pneuma-Ex+ for basal cell expansion, restoration of the expected epithelial morphology and gross composition can be achieved by using Thincert^TM^ membranes if differentiating in Pneuma-ALI or by using Pneuma-ALI-S instead of Pneuma-ALI on Transwell^TM^ membranes.

## Discussion and conclusions

Our study analyzes, at the single-cell level, the effect of 5 distinct cell culture media and 2 types of insert membranes for ALI differentiation of nasal and bronchial epithelial cells. All media allowed the detection of all major cell types present in the airway epithelium. We have shown that 2 cell culture media, Pneuma-ALI and H&H, generated ALI cultures with high MCC content. H&H contains half PneumaCult^TM^-ALI medium, minus the 100x supplement, which has a proprietary composition. We assumed that this supplement includes retinoic acid. The other half of this medium is composed of “Wu” medium which itself contains retinoic acid. According to the authors who have set-up this medium, it was supposed to create conditions that are more similar to *in vivo* conditions (34). However, except for some subtle differences, we found that cell identities and distributions were highly similar between Pneuma-ALI and H&H. On the contrary, the BEGM-ALI and Clancy media yielded much less MCCs. Thus, the choice of the culture conditions should be conditioned by the study to be performed. For cilia biology, Pneuma-ALI and H&H should be largely preferred as the MCC content was much higher in these conditions, with a neat deuterosomal cell cluster. However, if studying secretory cells, BEGM-ALI and Clancy media should be considered, taking into consideration the differences we found in secreted mucin expression. We have also noticed the rare occurrence of ionocytes, brush and pulmonary neuroendocrine cells, which were found in too little proportions to allow a comparison between media. Future investigations analyzing higher cell numbers should better identify the impact of differentiation media on the amounts and gene expression profiles of these cells.

We have used a label transfer method that utilizes reference similarity beyond marker genes alone to compare cells from the ALI model to *in vivo* airway cells. Given the very wide use of this culture model, such a comparison could be very useful. We have found some cell type and medium-dependent differences, mainly in basal, suprabasal, secretory and goblet cells, which are cell types showing continuous expression profiles, as opposed to MCCs, which have very distinct expression profiles and the largest number of marker genes. These data highlight the difficulty to annotate cell types based on automatic clustering followed by manual annotation and illustrate that using automatic annotations with label transfer that allow datasets to be compared to a reference in an unbiased manner tools could be a very useful complement.

Leung et al. have recently published a comparative study between Pneuma-ALI and BEGM-ALI (26). Their findings, in terms of multiciliated, goblet cell content and epithelial thickness, were very similar to ours. Saint-Cricq et al. have also compared Pneuma-ALI to UNC medium (University of Chapell Hill) which is quite similar to BEGM. They also obtained thicker epithelia with Pneuma-ALI and their differential gene expression analysis also showed increased expression of *IL33* and *TFF3* in Pneuma-ALI, and increased expression of ion transport genes in UNC medium, similarly to our findings in BEGM and Clancy media. Importantly, they showed that differentiation media can influence cystic fibrosis cell cultures responses to CFTR modulators (28). We add here that the choice of differentiation medium can also modify the expression of SARS-CoV-2 entry factors: ACE2 expression appeared lower in Pneuma-ALI, with a distinct glycosylation profile and *TMPRSS2* was restricted to MCCs in this medium, while it showed a wider expression in BEGM-ALI. Most studies have used Pneuma-ALI and found a higher SARS-CoV-2 tropism towards MCCs (16–21). However, V’kovski *et al.* and Johansen *et al.* used BEGM and found a higher tropism towards non-multiciliated cells (24, 25). Thus, medium selection might be a crucial parameter for viral infection studies.

We have extended our investigation to the stages occurring before differentiation. With the exception of few studies describing conditional reprogramming of epithelial cells (co-culture with irradiated fibroblast feeder cells and a Rho kinase inhibitor) (53, 54), or the use of dual SMAD signaling inhibition (55), these stages have been largely ignored in previous works using ALI cultures. Yet, proliferation conditions might influence cell fate by modifying identity and/or potency of basal cells which are the stem cells generating the entire epithelium upon setting-up the air-liquid interface. Here, scRNA-seq was instrumental to identify the precise cell composition of epithelial cell cultures during the initial propagation steps. We have shown that after a first stage on plastic flasks, cycling basal and basal cells are highly dominant in both bronchial and nasal cultures, but secretory, goblet, multiciliated and even endothelial cells were detected, with variations depending on the proliferation medium that was used. For nasal epithelial cells in Pneuma-Ex+ medium, we detected few proliferative goblet cells. These findings should be considered when using this medium and a more accurate characterization of these cells would be required. Interestingly, we found that gene expression profiles of basal cells were affected, with for instance, the upregulation of *SPDEF* by Pneuma-Ex+, which could influence the fate of these cells. In line with these findings, we found that proliferation in Pneuma-Ex+ favored the goblet cell lineage after differentiation.

Finally, we evaluated the effects of using an alternative differentiation medium (Pneuma-ALI-S) and alternative porous membranes (Thincerts^TM^) on epithelial morphology after proliferation in Pneuma-Ex+. We found that both restored the normal epithelial morphology which was lost with the use of Pneuma-Ex+ for all proliferation stages, followed by Pneuma-ALI differentiation. Since the composition of PneumaCult^TM^ products is not available, it is not possible to identify the differential cues that produced these effects. The use of alternative porous membranes has also been very informative and suggest that the pore density influences cell morphology. Thincerts^TM^ membranes having a lower pore density, we speculate that the flow of nutrient might be slower in these conditions and might account for the difference in epithelial morphology.

In conclusion, our study provides a reference for evaluating the influence of culture conditions on airway epithelial differentiation. Our scRNA-seq data could be further analyzed and could lead to the identification of other pathways regulating differentiation. Beyond the technical usefulness of our results, our study also identifies parameters that should be further explored to study cell fate and epithelial morphology.

