## Supplementary methods and figures for "Cell culture differentiation and proliferation conditions influence the *in vitro* regeneration of the human airway epithelium"

### Detailed Methods

#### Human Bronchial Epithelial Cells

Human Bronchial Epithelial Cells (HBECs) were obtained from human transplant donor lungs deemed unsuitable for transplantation and donated to medical research. The ethics committees of the institutions involved approved this study (CERC-SFCTCV-2018-5-6-9-8-32-DjXa). Cells were isolated from bronchial rings of 2 healthy donors, both non-smokers and with no chronic disease according to the family. Donors were a 62-year and 34-year old men. Primary human bronchial epithelial cells were isolated by protease digestion of bronchial rings. Briefly, after surgical removal specimens were rinsed in phosphate-buffered saline without $Ca^{2+}$ and $Mg^{2+}$ (PBS $^{-/-}$) to completely remove blood and mucus plugs. The epithelium on intact segments of bronchi was then dissociated with protease IV 0.1% (Sigma P5147) in PneumaCult$^{TM}$-Ex-Plus medium (STEMCELL Technologies) at 4°C for 16 hours. After digestion, the bronchial tube was rinsed with PBS $^{-/-}$ and dissociated cell clumps were strained through a 70-µm nylon mesh (Becton Dickinson). Cells were recovered and suspended in PneumaCult$^{TM}$-Ex-Plus medium containing 100 U/mL penicillin, 100 µg/mL streptomycin, 50 µg/mL gentamicin sulfate, and 2.5 µg/mL amphotericin B (all reagents from Gibco) and then plated on collagen I-coated 75 $cm^2$ flask. After 5 days of proliferation, after cells have reached about 80% confluence, cells were detached using trypsin-EDTA 0.05% (Gibco) for 5 min in 2 times, collected in inactivation buffer (DMEM supplemented with 10% heat-inactivated fetal bovine serum (FBS), 100 U/mL penicillin, 100 µg/mL streptomycin, 50 µg/mL gentamicin sulfate, and 2.5 µg/mL amphotericin B), centrifuged at 150 g for 5 min and resuspended in 10 mL HBSS with 0,04% BSA. Cells were filtered through a 15 µm cell strainer (Pluriselect) to remove clumps and centrifuged. Cells were counted with Countess™ automated cell counter (Thermo Fisher Scientific). One part of the cells was used to capture 1000 cells by the Chromium device (10X genomics) to perform 3' single-cell RNA-seq experiment. The rest of the cells were split in 4, centrifuged and resuspended in 4 distinct proliferation media: (1) PneumaCult™ Ex-Plus medium, (2) Wu medium (Table 1), (3) BEGM™ (Lonza, Bronchial Epithelial Growth Medium, prepared as indicated by the manufacturer, but omitting gentamycin) and (4) Clancy medium (Table 2). All proliferation media contained 100 U/mL penicillin, 100 µg/mL streptomycin, 50 µg/mL gentamicin sulfate, and 2.5 µg/mL amphotericin B. Cells were seeded on human placenta collagen IV-coated Transwell™ permeable supports (reference 3470, 6.5 mm diameter; 0.4 µm pore size; Corning) in the apical part with a density of ~50 000 cells per Transwell™ and with medium in the basal part. The remaining cells were used to perform single-cell RNA-seq (scRNA-seq). Medium was replaced 48h after seeding, and once the cells have reached confluence (~5 days after seeding), the medium was removed from the apical side of the Transwell™ and the basal medium, replaced by differentiation media: (1) PneumaCult™-ALI (STEMCELL Technologies), (2) Half & Half, (3) BEGM™ -ALI medium, (4) Clancy medium (Table 2). BEGM™-ALI was prepared by adding BEGM™ SingleQuot™ (Lonza) to BEBM™:DMEM (50:50) but omitting the gentamycin and replacing retinoic acid so as to obtain a final concentration of 30 nM (Sigma Aldrich reference R2625). Half & Half medium was Wu medium:Pneumacult-ALI medium (50:50) with all PneumaCult™-ALI supplements but omitting the 100x supplement (1). All differentiation media contained 100 U/mL penicillin, 100 µg/mL streptomycin and 50 µg/mL gentamicin sulfate. The day the air-liquid interface was applied, and media were replaced by differentiation media were named “ALI0” for “Air-Liquid 0”. Cells from each proliferation condition were dissociated (see below) to perform a scRNA-seq analysis. Culture medium was changed every two days. Cells were harvested at fully-differentiated state after 28 days (ALI28).

**Table 1: Composition of Wu medium** (2, 3)

| Product | Supplier | Reference | Final concentration |
|---|---|---|---|
| Hepes | Gibco | 15630080 | 15 mM |
| MgCl2 | Sigma | M1028 | 0.3 mM |
| MgSO4 | Sigma | M3409 | 0.4 mM |
| CaCl2 | Alfa Aesar | J62905-AP | 1.05 mM |
| Bovine pituitary extract | Sigma | P1476 | 15 µg/mL |
| All trans retinoic acid | Sigma | R2625 | 30 nM |
| Insulin | Sigma | I6634 | 5 µg/mL |
| Transferrin | Sigma | T0665 | 5 µg/mL |
| EGF | Corning | 354052 | 10 ng/mL |
| Cholera toxin | Sigma | C8052 | 20 ng/mL |
| BSA | Sigma | | 5 mg/mL |
| Dexamethasone | Sigma | D4902 | 0.5 µM |

**Table 2: Composition of Clancy medium** (2)

| Product | Supplier | Reference | Final concentration |
|---|---|---|---|
| Hepes | Gibco | 15630080 | 15 mM |
| Ultroser-G | Pall | | 0.02 |
| Fetal Clone II | Hyclone | SH 30066-03 | 0.02 |
| Bovine Brain extract | Lonza | CC-4092 | 0.0 |

| | | | |
|---|---|---|---|
| Alltrans retinoic acid | Sigma | R2625 | 10 nM |
| Insulin | Sigma | I6634 | 2.5 µg/mL |
| Transferrin | Sigma | T0665 | 5 µg/mL |
| Hydrocortisone | Sigma | H0396 | 20 nM |
| Triiodothyronine | Sigma | T6397 | 500 nM |
| Epinephrine | Sigma | E4250 | 1.5 µM |
| Phosphoethanolamine | Sigma | P0503 | 250 nM |
| Ethanolamine | Sigma | E0135 | 250 nM |

#### Human Nasal Epithelial Cells

Human nasal epithelial cells (HNECs) were obtained from inferior turbinates that were resected from patients who underwent surgical intervention for nasal obstruction or septoplasty (kindly provided by Professor Castillo, Pasteur Hospital, Nice, France). The use of human tissues was authorized by the bioethical law 94–654 of the French Public Health Code after written consent from the patients. After surgery, nasal inferior turbinates were immediately immersed in cold $Ca^{2+}$/$Mg^{2+}$-free HBSS supplemented with 25 mM HEPES, 100 U/mL penicillin, 100 µg/mL streptomycin, 50 µg/mL gentamicin sulfate, and 2.5 µg/mL amphotericin B (all reagents from Gibco). After repeated washes with ice-cold supplemented HBSS, turbinates were immersed in supplemented HBSS containing 0.1% protease XIV from Streptomyces griseus overnight at 4°C for epithelial digestion. Gentle agitation allowed to collect detached cells in inactivation buffer (DMEM supplemented with 10% FBS, 100 U/mL penicillin, 100 µg/mL streptomycin, 50 µg/mL gentamicin sulfate, and 2.5 µg/mL amphotericin B). After centrifugation at 150 g for 5 min, cells were resuspended in inactivation buffer, passed through a 21-G needle for additional mechanical dissociation, split into several tubes depending on the number of different proliferation media used and centrifuged again. Cells were then resuspended in different proliferation media depending on the experiment, as indicated in the figure legends, plated (~20 000 cells per cm²) on 75 cm²-flasks coated with rat-tail collagen I (Sigma-Aldrich C3867), and incubated in a humidified atmosphere of 5% $CO_2$ at 37°C. After 24h, cells were rinsed with PBS to remove erythrocytes and debris from the culture and immerged again in proliferation medium. After 4 to 5 days of culture, cells reached 70% confluence. Cells were detached using trypsin-EDTA 0.05% for 5 to 10 min, collected in inactivation buffer as described above, centrifuged at 150 g for 5 min and resuspended in their respective proliferation media. Cells were seeded on human placenta collagen IV-coated Transwell™ permeable supports (reference 3470, 6.5 mm diameter; 0.4 µm pore size; Corning) in the apical part with a density of ~30,000 cells per Transwell™ (or otherwise specified) and with medium in the basal part. Once the cells reached confluence (~5 days after seeding), the medium was removed from the apical side of the Transwell™. The basal medium was replaced by differentiation medium depending on the experiment, as indicated in the figure legends, to create Air-Liquid Interface (ALI) cultures as previously described. This day corresponded to the differentiating step at day 0 (ALI0). Culture medium was changed every other day. All media were supplemented with 100 U/mL penicillin, 100 µg/mL streptomycin, 50 µg/mL gentamicin sulfate, and 2.5 µg/mL amphotericin B (all from Gibco).

To assess the effects of proliferation media (Supplementary Figure 1C), dissociated cells were resuspended in either Bronchial Epithelium Basal Medium (BEBM™, Lonza) supplemented with BEGM™ SingleQuot™ Kit supplements (Lonza) (gentamycin from the kit was replaced with gentamycin from Gibco at 50µg/ml) or in PneumaCult™-Ex Plus, and spent two passages on T75 flasks. Either BEGM™-ALI (as described above) or PneumaCult-ALI™ were used as differentiation medium.

For the single-cell RNA-seq analysis of proliferating nasal epithelial cells (Figure 3D), cells isolated as described above, passaged once in BEBM™ and cryopreserved in CryoStor® CS10 (STEMCELL Technologies). They were then thawed in BEBM™, split in two tubes, centrifuged at 150 g and resuspended in PneumaCult™-Ex or PneumaCult™-Ex Plus. At 70% confluence cells were isolated for single-cell RNA sequencing as described below.

For the comparison of media alternatives experiment (Figure 6), HNECs were isolated and expanded in PneumaCult™-Ex supplemented extemporaneously with Y27632 at 10 µM (STEMCELL Technologies), or PneumaCult™-Ex+ as described above. Cells were seeded at ~50 000 cells per collagen IV–coated Transwell™ membrane. Either PneumaCult™-ALI or PneumaCult™-ALI-S (STEMCELL Technologies) were used as differentiation medium.

For the comparison of commercial membranes experiment (Figure 7), HNECs were isolated and expanded in PneumaCult™-Ex as described above. Cells were seeded at ~50 000 cells per collagen IV –coated membrane, either Transwell™ or Thincert™ (Greiner 662641). At confluence the basal medium was replaced by differentiation medium: either PneumaCult™-ALI or Promocell-ALI to create Air-Liquid Interface (ALI) cultures. Promocell-ALI was obtained by mixing 50:50 Airway Epithelial Cell Growth Medium (Promocell):DMEM and adding all media supplements provided with the PromoCell® Airway Epithelial Cell Growth Medium, replacing retinoic acid from the kit in order to reach a final concentration of 30 nM (Sigma Aldrich reference R2625), similarly to BEGM-ALI.

#### Paraffin embedding and H&E staining

Fully differentiated epithelia on Transwells™ or Thincerts™ were fixed in paraformaldehyde 4% at room temperature for 15 min, then extensively rinsed with phosphate-buffered saline (PBS). Each membrane was cut with a razor blade, then dehydrated in successive baths of xylene and ethanol and embedded in paraffin. Cutting of paraffin sections was performed in a rotary microtome MICROM HM 340E (Thermo Fisher Scientific). Before immunofluorescence staining, deparaffinization and antigen retrieval process were carried out using citrate buffer at pH6, then permeabilized with 0.5% Triton X-100 in PBS and blocked with 3% BSA in PBS for 30 min. Sections were incubated with primary antibodies diluted in PBS-BSA 3% at 4°C overnight. Incubation with secondary antibodies was carried out during 1h at room temperature after 3 PBS washes of 5 minutes. Nuclei were stained with 4,6-diamidino-2-phenylindole (DAPI).

For hematoxylin and eosin staining, sections were deparaffinized and immersed in hematoxylin (Harris) for 3 min 30 s followed by immersion in eosin for one minute. Slides were then thoroughly rinsed, dehydrated in ethanol and xylene and mounted with Eukitt® (ORSAtec GmbH).

#### Immunostainings

Fully differentiated epithelia on Transwells™ or Thincerts™ were fixed in 4% paraformaldehyde for 15 min at room temperature. After 3 PBS washes, cells were permeabilized with 0.5% Triton X-100 in PBS for 5 min at room temperature. Then, cell layers were blocked with 3% BSA in PBS for 45 min at room temperature. Cells were incubated with primary antibodies, diluted at the required concentration in 3% BSA in PBS (Table 3) at 4°C overnight. After 3 washes of 5 min with PBS, incubation with secondary antibody (Table 4), diluted in 3% BSA in PBS, was carried out during 1 hour at room temperature and protected from light. Nuclei were stained with 4,6-diamidino-2-phenylindole (DAPI), diluted at 1/1000 in the secondary antibody solution. Phalloidin-Alexa Fluor™ 594 (Thermo Fisher A12381, 1/250), was used when indicated. It was added to the secondary antibody solution. After 3 washes in PBS, Transwells™ membranes were cut with a razor blade and mounted with ProLong™ Diamond (Invitrogen™) mounting medium. Images were acquired using either Zeiss Axioplan2, Olympus Fv10i, Leica SP5 or SP8 confocal imaging systems.

**Table 3: References and dilutions of the primary antibodies used in the study**

| **Target** | **Clone** | **Supplier** | **Catalog number** | **Dilution** | **Host species/ Isotype** |
|---|---|---|---|---|---|
| Acetylated Alpha-tubulin | 6B11 | Merck | T7451 | 1/1000 | Mouse / IgG2b |
| MUC5AC | 45M1 | Abnova | MAB11324 | 1/500 | Mouse / IgG1 |
| MUC5B | 6F10-E4 | Cancertools | EU-MUC5Ba | Mix 1:1 and use 1/500 | Mouse / IgG1 |
| MUC5B | 6F10-E4 | Cancertools | EU-MUC5Ba | | Mouse / IgG1 |
| KRT5 | Poly19055 | Ozyme | BLE905501 | 1/2000 | Rabbit |
| ACE2 | Polyclonal | R&D systems | AF933 | 1/400 | Goat |
| FOXJ1 | 2A5 | eBioscience | 14-9965-82 | 1/200 | Mouse / IgG1 |
| GAPDH | 1E6D9 | Proteintech | 60004-1 | 1/20000 | Mouse / IgG2b |
| Tubulin | 1E4C11 | Proteintech | 66031-1 | 1/1000 | Mouse / IgG2b |

**Table 4: References and dilutions of the secondary antibodies used in the study**

| Antibody name | Supplier | Catalog number | Dilution | Application |
|---|---|---|---|---|
| Anti-mouse IgG1 - Alexa Fluor 488 | Invitrogen | A-21121 | 1/500 | Immunofluorescence |
| Anti-mouse IgG2b - Alexa Fluor 647 | Invitrogen | A-21242 | 1/500 | Immunofluorescence |
| Anti-rabbit- Alexa Fluor 647 | Invitrogen | A-11037 | 1/500 | Immunofluorescence |
| Anti-Goat -HRP | Jackson Immunoresearch | 305 036 045 | 1/5000 | Western blot |
| Anti-Rabbit -HRP | Dako | P0448 | 1/5000 | Western blot |
| Anti-mouse -HRP | Dako | P0447 | 1/5000 | Western blot |

#### SP5/SP8 confocal acquisitions

Confocal images were acquired using either a Leica SP8 STED 3X (Leica Microsystems, Nanterre) at 700Hz or a Leica TCS SP5 at 200Hz. Confocal images were obtained through a 63x/1.4 NA Oil objective using the LAS X software (Leica Microsystems, Nanterre), by a 405, 488 and 561 nm laser excitation respectively.

#### Gene expression analysis by real-time qPCR

RNAs were extracted from proliferating basal cells or differentiated ALI epithelium with the miRNeasy micro kit (Qiagen) and then retro-transcribed with the RT High-capacity kit (Thermofisher Scientific). Quantitative PCR was performed using LightCycler® 480 SYBR Green I Master mix and Light Cycler 480 real-time PCR machine (Roche Applied Science). Expression levels of transcripts were evaluated using the comparative CT method (2-deltaCT). Transcript levels of TBP were used for sample normalization. Primers sequences are detailed in Table 5. Results are either 2-deltaCT (FOXJ1-TBP and MUC5AC-TBP) for Figure 4 or log2-transformed fold changes of normalized 2-deltaCT (SPDEF-TBP) for Supplementary Figure 10.

**Table 5: Sequences for the PCR primers used in the study**

| **Target gene** | **Primer sequences** |
|---|---|
| *SPDEF* | 5’- GTCTGACTTCCTCCCAGCAC - 3’<br>5’- CTTGGAGGACTGGGTCTGTG - 3’ |
| *FOXJ1* | 5’- TGGATCACGGACAACTTCTG - 3’<br>5’- GAGGCACTTTGATGAAGCAC - 3’ |
| *MUC5AC* | 5’- CCAAATACGCCAACAAGACC- 3’<br>5’- ATTCCATGGGTGTCAGCTTG- 3’ |
| *MUC5B* | 5’- GCTCCAAGGCCATCAAGCT- 3’<br>5’- GGTCTCGATGACCAGGAAGATC- [illegible] |
| *ACE2* | 5’- TGGGACTCTGCCATTTACTTAC - 3’<br>[illegible] |
| *TBP* | 5’- ACGCCAGCTTCGGAGAGTTC - 3’<br>5’- CAAACCGCTTGGGATTATATTCG [illegible] |

#### Western blot experiments

Primary HNECs from differentiated ALI epithelia were harvested by scraping in anti-protease-supplemented Ripa lysis buffer (Thermo Scientific Pierce), cleared by centrifugation and ultrasonicated. Protein concentration was determined using the Bio-Rad Protein Assay (Bio-Rad) and equivalent amounts of protein were resolved on SDS polyacrylamide gels using Novex™ NuPAGE™ SDS/PAGE Gel System following the manufacturer’s instructions. Proteins were transferred to PVDF membranes and analyzed by immunoblotting with appropriate primary antibodies and HRP-conjugated secondary antibodies (1:5000, Dako, Agilent Technologies) according to the manufacturer’s instructions. Primary antibodies and dilution used are indicated in Table 3.

Immunoreactive bands were detected using immobilon ECL kit (Merck Millipore, Billerica, MA, USA) on a LAS-3000 imager (Fujifilm France, BOIS D'ARCY, France).

#### Cell dissociation for single-cell RNA-seq

To perform single-cell analysis, cells grown on Transwells™ were harvested by incubating 15 min at 37°C (ALI0 time point) with trypsin EDTA or for 4-6 hours at 4°C with 0.1% protease type XIV *Streptomyces griseus* in HBSS for the ALI28 time point until dissociation was clearly visible. Then, cells were gently detached from the Transwell™ by pipetting and transferred into a microtube. An excess of HBSS containing 2% BSA was added. Cells were centrifuged at 150 g for 5 min and resuspended in 500 µL of supplemented HBSS containing 2% BSA, centrifuged again at 150 g for 5 min and resuspended in 500 µL HBSS with 10% FBS and dissociated mechanically 4 times through a 26G syringe. Finally, cell suspensions were filtered through a 40 µm porosity Flowmi™ Cell Strainer (Bel-Art), centrifuged at 150 for 5 min and resuspended in 500 µL of HBSS. Cell concentration and viability were measured with a Countess™ automated cell counter (Thermo Fisher Scientific), after incubation with NucGreen to detect dead cells (Thermo Fisher scientific). All steps, except trypsin incubation were performed on ice.

For scRNA-seq after the initial cell propagation on plastic flasks, cells were harvested by incubating for 5 min at 37°C with trypsin-EDTA, inactivated with an excess of HBSS 2% BSA and processed as described above. Cell concentration was adjusted to 300 cells/µL in HBSS for the cell capture by 10X genomics device, in order to capture 1750 cells. For ALI0 and ALI28, cell hashing was performed (see below).

#### Cell Hashing for single-cell RNA-seq

Cell concentration of each cell suspension (4 conditions for ALI0 and ALI28 time point) was determined, and 500 000 cells of each were centrifuged and resuspended in 100 µL of HBSS containing 2% BSA. Each cell suspension was then incubated with 0.5 µg of a distinct hashtag antibody (TotalSeq™-A0251, A0252, A0253 and A0254, BioLegend). Each cell suspension was washed twice with HBSS, and cell concentration was measured again. Cells from each of the 4 conditions were pooled in order to obtain a final cell suspension containing identical cell concentrations of each condition. We aimed at reaching a total concentration of 700 cells/µL in HBSS for the cell capture by 10X genomics Chromium, in order to capture 10 000 cells i.e. 2500 cells per condition.

#### Single-cell RNA-seq

We followed the manufacturer's protocol (Chromium™ Single Cell 3' Reagent Kits, v2 for HBEC dataset and v3 Chemistry for HNEC dataset) to obtain single cell 3' libraries for Illumina sequencing. For HBEC dataset the libraries were sequenced with a NextSeq 500/550 High Output v2 kit (75 cycles): Read 1 had a length of 26 bases that included the cell barcode and the UMI; Read 2 had a length of 57 bases that contained the cDNA insert; Index reads for sample index of 8 bases. For HNEC dataset the library was sequenced with a NextSeq 2000 P3 Reagents (100 cycles) in paired-end mode at length of 28 bases for R1 and 90 bases for R2 with 2 index reads of 10 bases. Raw sequencing data were processed using the 10x Cell Ranger count pipeline (v6.0.0 for HBEC dataset and v7.0.0 for HNEC dataset), with default parameters and aligned to the GRCh38 human reference genome (Gencode Release 38). For cell hashing experiments (ALI0 and ALI28 from HBEC dataset), we generated antibody count matrices from fastq files using CITE-seq-Count (v1.4.2) (https://github.com/Hoohm/CITE-seq-Count) with the following command: CITE-seq-Count -R1 X_R1_001.fastq.gz -R2 X_R2_001.fastq.gz -T 4 -t HTO.barcodes.biolegend.human.csv -wl barcodes_X.tsv --output hashtag_count_X-cbf 1 -cbl 16 -umif 17 -umil 26 -cells 10000.

#### Single-cell data analysis

*Preprocessing, integration, normalization and clustering.* Individual dataset analysis was performed using Seurat standard analysis pipeline v4.3.0 (4). First, for cell hashing samples (ALI0, ALI28), we used the HTODemux function of Seurat (default parameters) to assign antibody tags for each cell based on the antibody count matrices. Then, for all samples, cells were filtered out based on number of expressed features, dropout percentage, library size and mitochondrial gene percentage. Thresholds were selected by visually inspecting violin plots in order to remove the most extreme outliers (for samples details and filtering see Supplementary Table 7). We performed integration with either Seurat integration or scVI integration (latent space of 50 dimensions) (v0.14.6) (5). We constructed a k-nearest neighbor graph with default parameters. The Uniform Manifold Approximation and Projection (UMAP) representation was used for visualization of integrations into two dimensions. Cell-level normalization was performed using NormalizeData function. Highly variable genes were selected for following analysis based on their expression level and variance. Clustering was first performed with default parameter using Seurat function. Differential analysis was again performed using Seurat FindAllMarkers and FindMarkers functions based on non-parametric Wilcoxon rank sum test.

*Cell Type Annotation.* We used FindAllMarkers function to find gene markers for each cluster found previously. We then annotated on the basis of these gene markers and by visualizing the expression of our gene markers (Supplementary Table 1).

*Analysis of cell type proportions.* We used the propeller function (with asin transformation) from speckle package (v0.0.3) (6, 7) to compare cell type proportion between the different media. Supplementary Table 2 shows the detailed cell type proportions by medium.

*Differential Expression.* We performed differential expression analysis by using the run_de function from Libra package (v1.0.0) (https://github.com/neurorestore/Libra) and using pseudobulk method with edgeR. We compared each medium 2-by-2 at ALI28. Supplementary Table 3 shows the differential gene expression output.

*Label transfer to the Human Lung Cell Atlas (HLCA)* (8)*.*

We utilized the same method outlined by the authors of the UCE manuscript with minor extensions (9). We first embedded both our generated media datasets and the human lung cell atlas core scRNA-seq data in the UCE embedding space using the 33-layer model provided by the authors. We then calculated a cell type centroid for each unique cell type label within the HCA lung core dataset based on the specified annotation level, by calculating the mean of the cell vectors for each cell in that cell type class. We then classified every cell in the media datasets (the query datasets) based on the nearest centroid using cosine distance. This is how we defined the label transfer, and for the transferred or “predicted” labels, we can define confusion matrices for each media dataset. We then derived an accuracy score, an F1 score, and also calculate the average distance to centroid for all points within each cell type category. The code used to implement this methodology are available at the following GitHub repository:

https://github.com/Eamonmca/Single-Cell-Fuzzy-Labels

*CellChat analysis.* We computed individual CellChat analysis (v1.4.0) (10) for each medium at ALI28. We followed the tutorial of CellChat with default parameters (https://github.com/sqjin/CellChat).

*Data availability.* Single-cell RNA-seq data is available under GEO accession GSE243045 and EGA accession EGAD00001011362.

*Supplementary references.*

6. Phipson B, Sim CB, Porrello E, Hewitt AW, Powell J. propeller : testing for differences in cell type proportions in single cell data. 2021;

7. Phipson B, Sim CB, Porrello ER, Hewitt AW, Powell J, Oshlack A. propeller: testing for differences in cell type proportions in single cell data. In: Mathelier A, editor. *Bioinformatics* 2022;38:4720–4726.

8. Sikkema L, Ramírez-Suástegui C, Strobl DC, Gillett TE, Zappia L, Madissoon E, *et al.* An integrated cell atlas of the lung in health and disease. *Nat Med* 2023;29:1563–1577.

9. Rosen Y, Brbić M, Roohani Y, Swanson K, Li Z, Leskovec J. Toward universal cell embeddings: integrating single-cell RNA-seq datasets across species with SATURN. *Nat Methods* 2024;doi:10.1038/s41592-024-02191-z.

10. Jin S, Guerrero-Juarez CF, Zhang L, Chang I, Ramos R, Kuan C-H, *et al.* Inference and analysis of cell-cell communication using CellChat. *Nat Commun* 2021;12:1088.

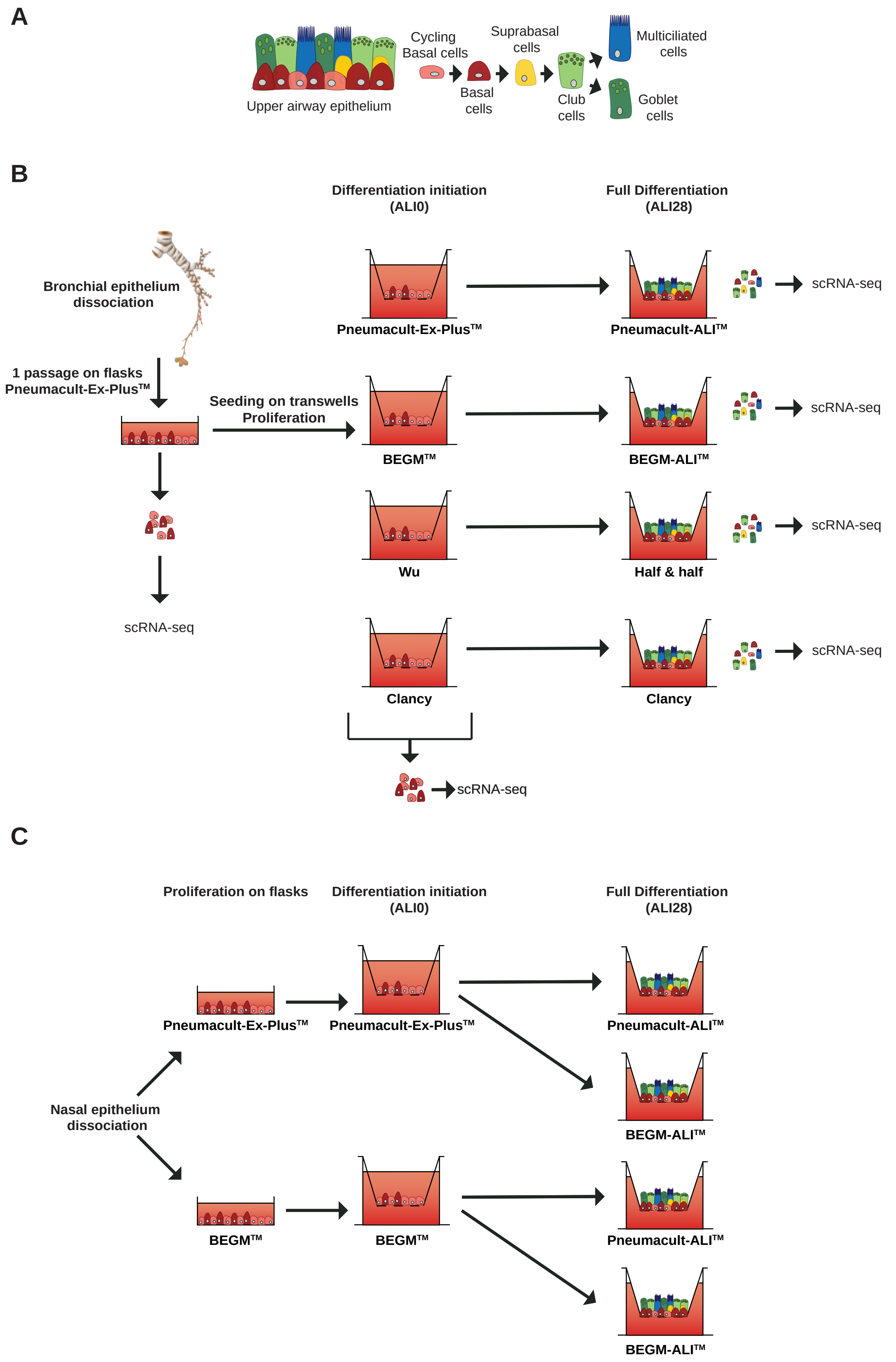


**Supplementary Figure 1: Overview of the experimental strategy.**
**(A)** Model of upper airway epithelium, based on six major types of epithelial cells, with consensus lineage hierarchy. **(B)** scRNA-seq experimental design. Bronchial airway epithelial regeneration was performed in 4 distinct cell culture media: after a first cell expansion on collagen-coated plastic flasks in Pneuma-Ex+ medium, human bronchial epithelial cells (HBECs) were dissociated and seeded on collagen-coated Transwells™ membranes in 4 distinct proliferation media. Differentiation was induced by the transition to an air-liquid interface (ALI) and the use of 4 distinct differentiation media. Cells were analyzed after the first expansion in Pneuma-Ex+, at the onset of differentiation (ALI0) and at full differentiation (ALI28). Cells were isolated from bronchi of two independent healthy donors. **(C)** Experimental design to assess the effects of proliferation media on human nasal epithelial cells (HNECs). After cell expansion on collagen-coated plastic flasks in either Pneuma-Ex+ from BEGM, each HNEC culture was dissociated and seeded on collagen-coated Transwells™ membranes in 2 distinct proliferation media.

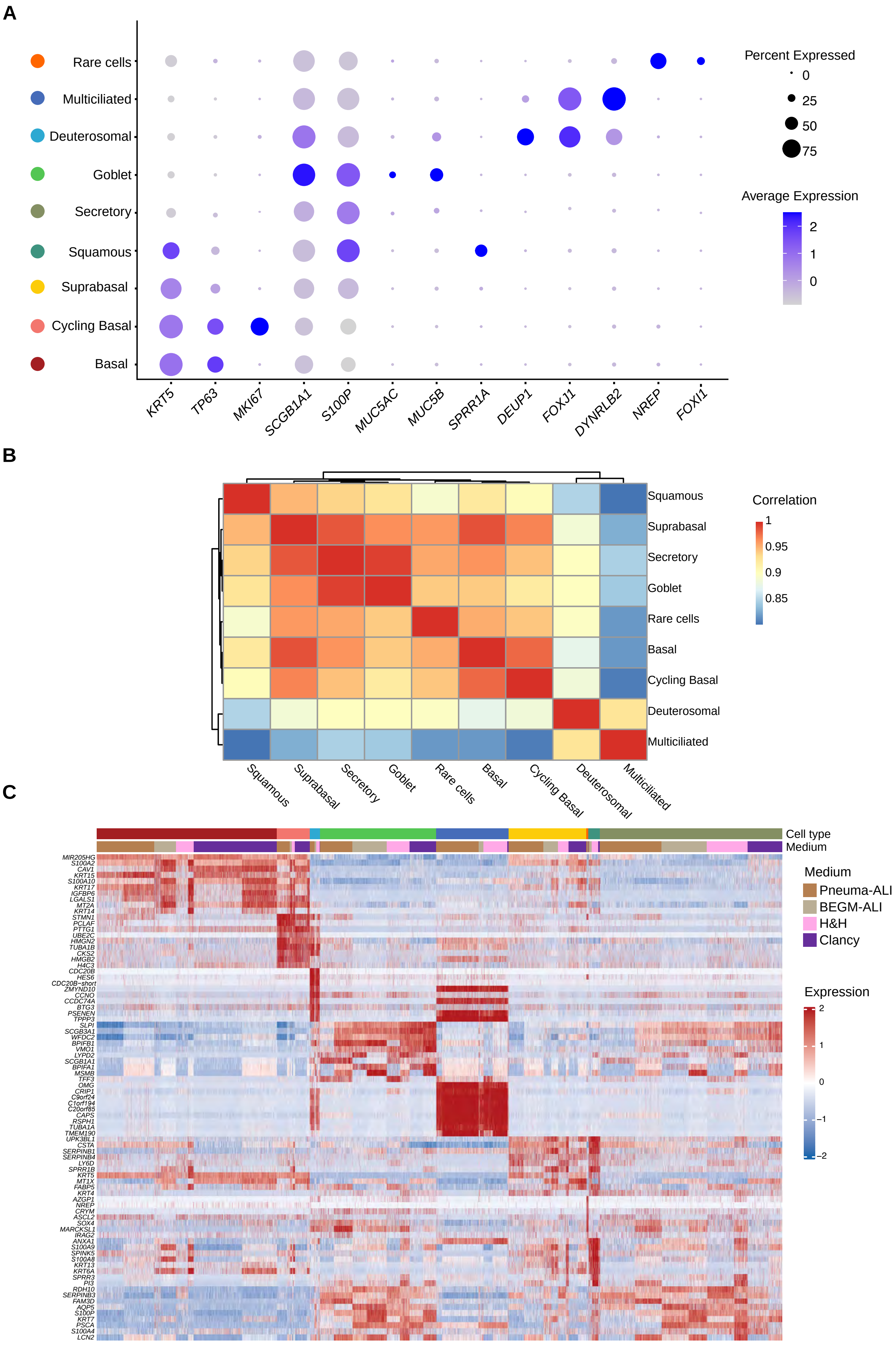


**Supplementary Figure 2: scRNA-seq gene expression profiles of cell clusters at ALI28 of bronchial epithelial cell differentiation.**
**(A)** Dotplot showing the expression of main marker genes that are usually selected for cell typing, in the integrated ALI28 dataset. **(B)** Gene expression profile correlations between cell clusters of the integrated ALI28 dataset. **(C)** Heatmap showing the top 10 specific genes for each cell, for each medium.

**A**

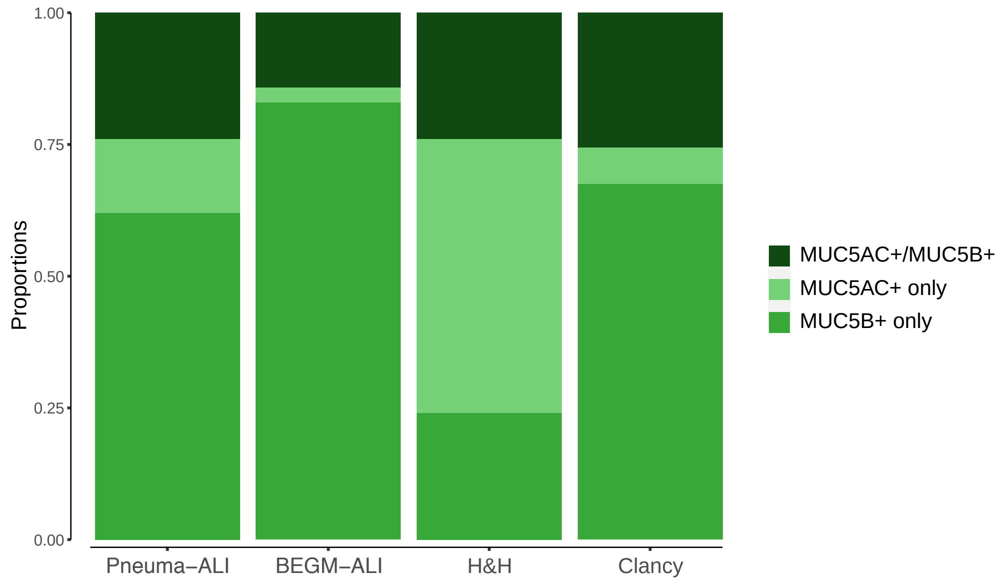


**B**

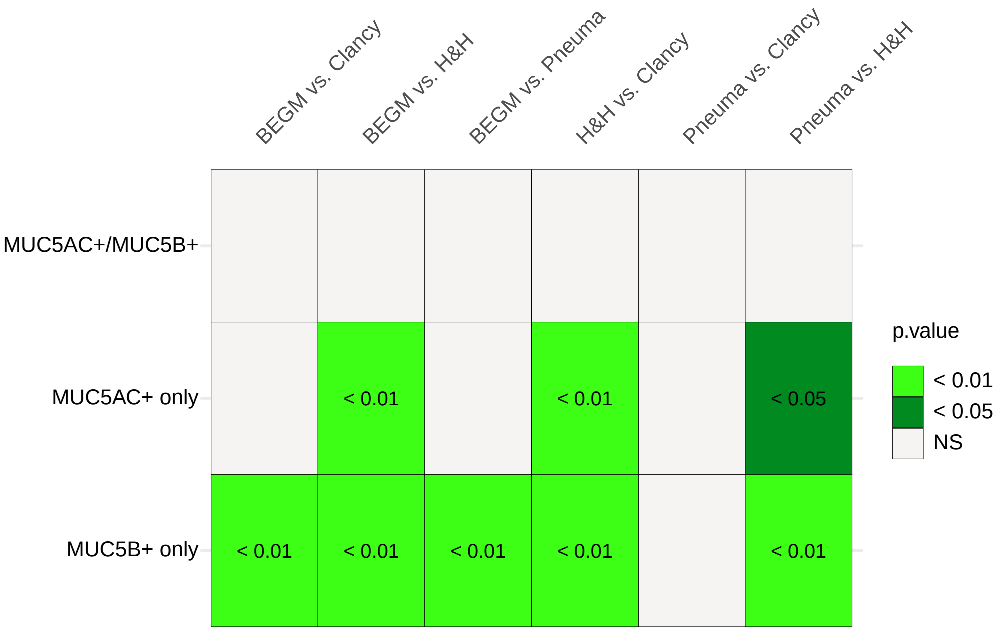


**Supplementary Figure 3: Proportions of MUC5AC+ only, MUC5B+ only or MUC5AC+/MUC5B+ goblet cells in each differentiation medium, according to scRNA-seq.**
**(A)** Stacked bar plot showing proportions of MUC5AC+ and/or MUC5B+ goblet cells. MUC5AC and or MUC5B+ cells were identified as positive if expression was above a cutoff that was defined as the 98th percentile of expression in non-goblet cells.
**(B)** p-value of t-test by propeller package, using the asin transformation, for each 2-to-2 medium comparison of MUC5AC+ or MUC5B+ or MUC5AC+/MUC5B+ proportions.

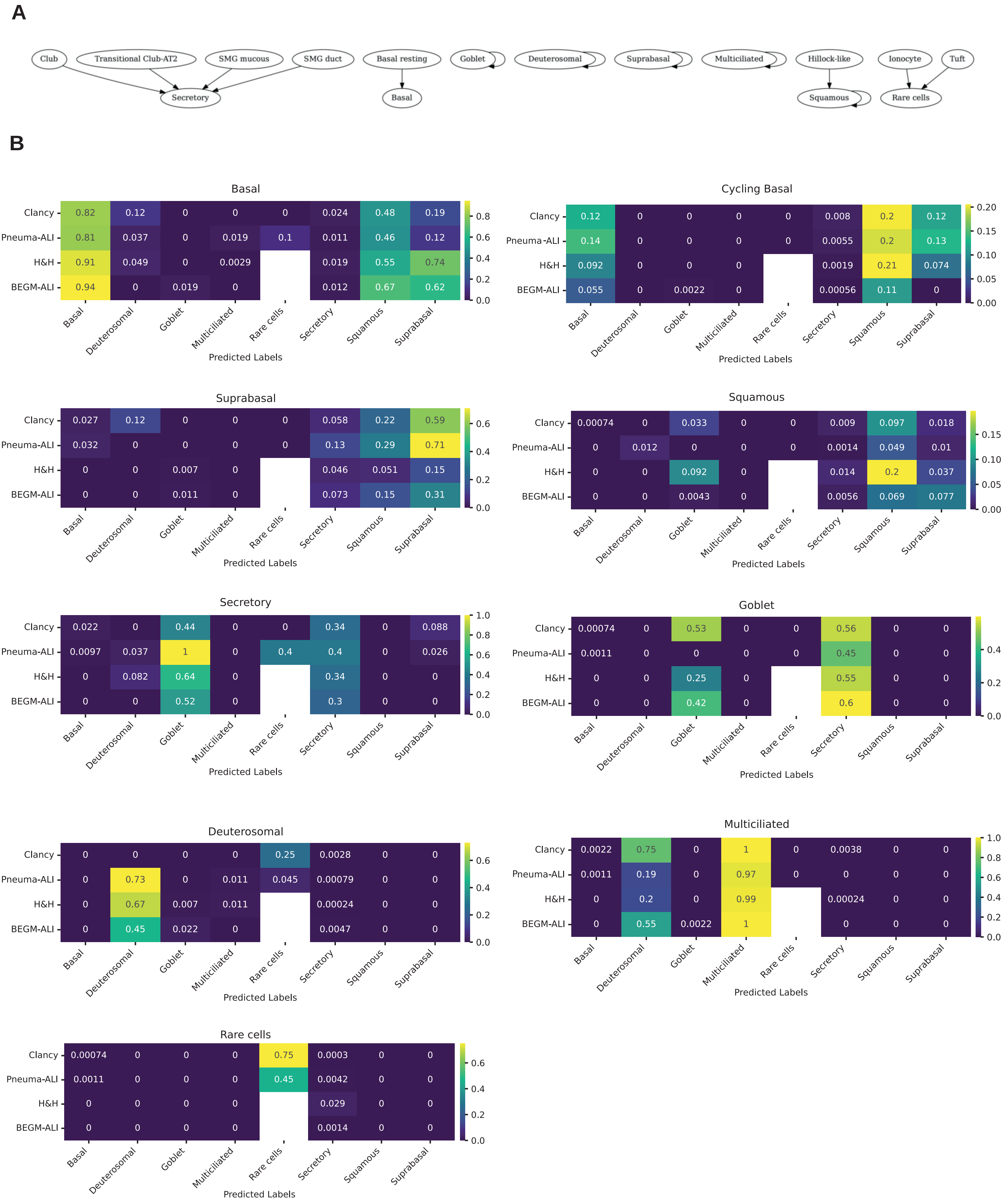


**Supplementary Figure 4: Label transfer from single-cell RNAseq data at ALI 28 in 4 differentiation media towards the Human Lung Cell Atlas (HLCA) reference.**
**(A)** Label harmonization scheme between the query (ALI28 dataset) and reference (HLCA). **(B)** Heatmap derived from confusion matrices organized by cell type from the ALI28 dataset, transferred to the HLCA reference, showing prediction proportions for each cell type. Maximal score is 1.

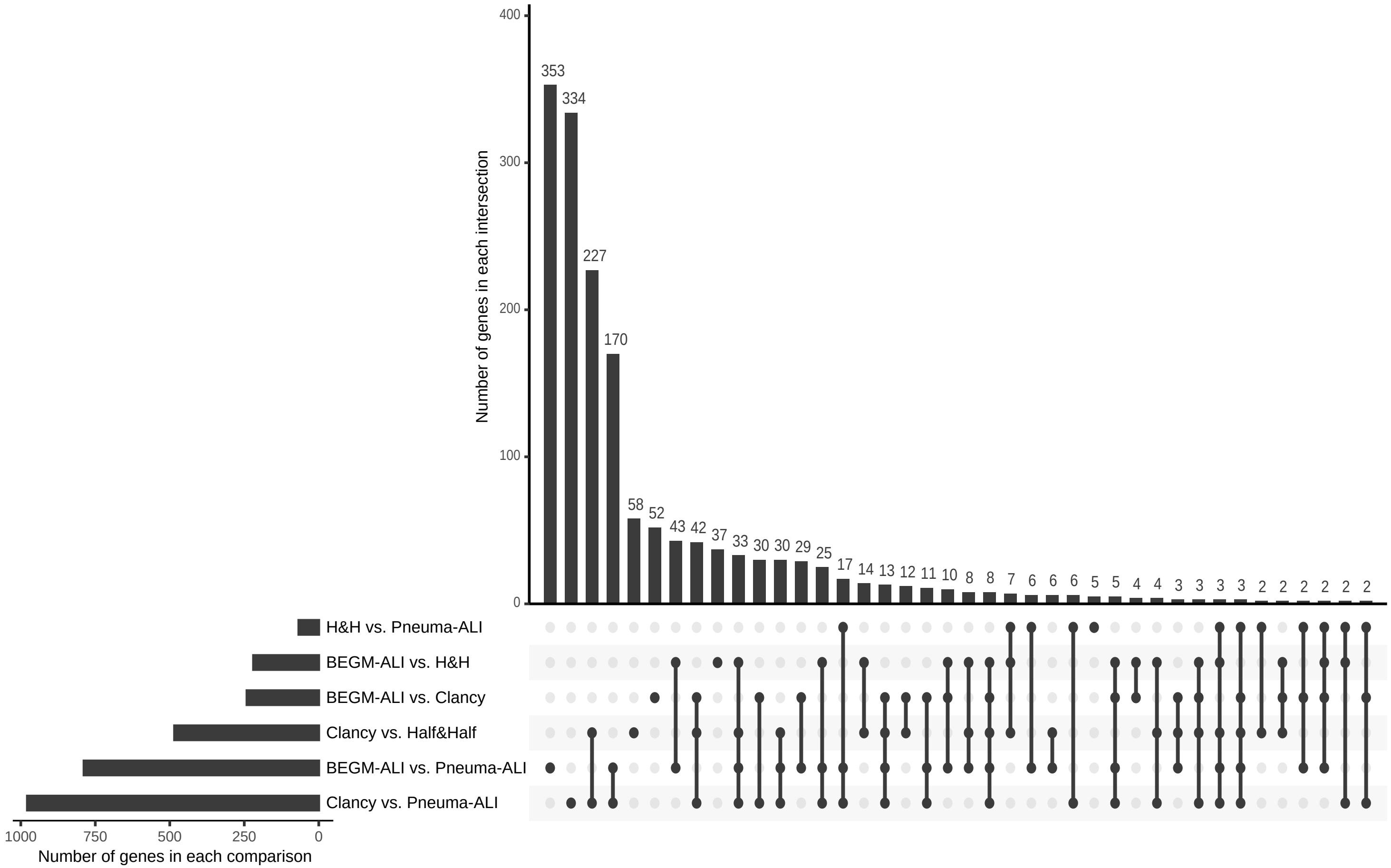


**Supplementary Figure 5: Effect of differentiation medium on gene expression at ALI28 of bronchial epithelial cell differentiation.**
Upset plot showing number of differentially expressed genes obtained by a pseudobulk strategy comparing media by pairs (Bar plot showing number of genes in each comparison). Genes differentially expressed in at least one cell type were included. The upset plots show number of genes detected in each intersection of media comparison. A single dot with no joining bar between lines shows comparisons for which genes were detected in a unique manner.

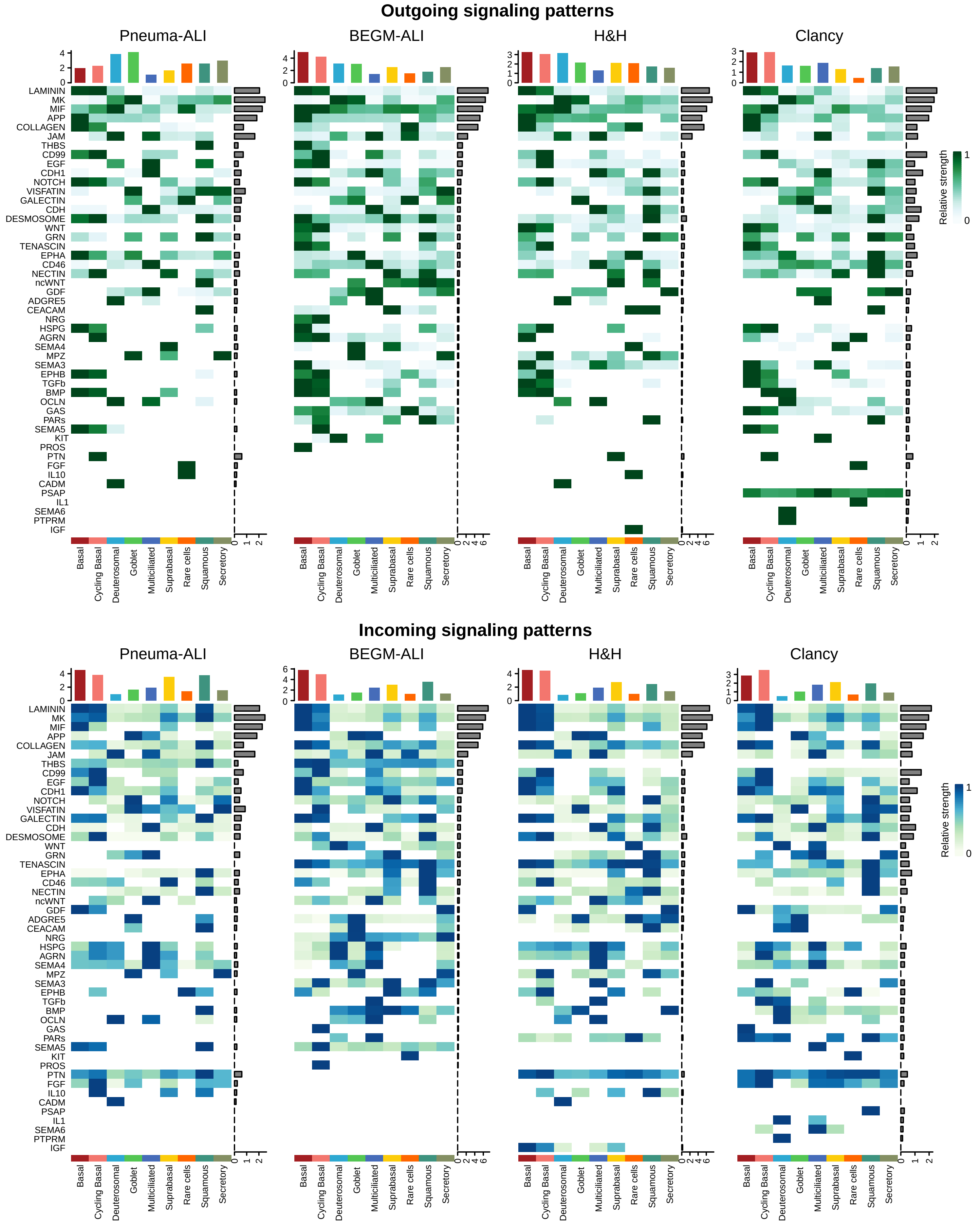


**Supplementary Figure 6: CellChat heatmaps of the significant outgoing and incoming signaling patterns in each differentiation media at ALI28 of bronchial epithelial cell differentiation.** Bars represent the outgoing/incoming overall potential on each cluster (top) and pathway (right).

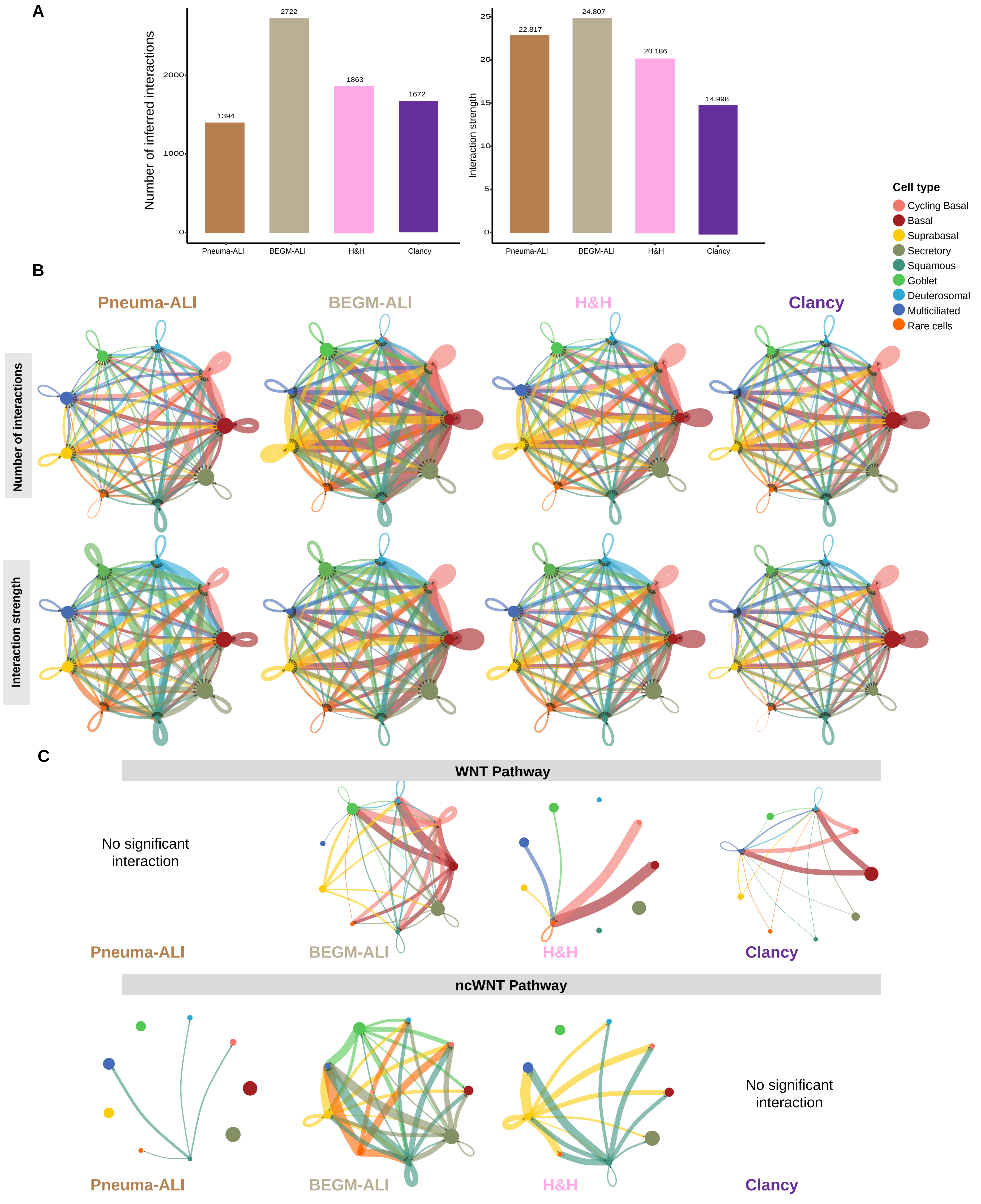


**Supplementary Figure 7: Ligand-receptor interactions at ALI28 of bronchial epithelial cell differentiation.**
**(A) Bar plots showing number of interactions (left) and interaction strength (right) for each differentiation medium. (B)** Circle plots showing communication probabilities (number of interactions and weights or interaction strength as indicated) of significant inferred ligand-receptor pairs between any pair of two cell populations. The edge width is proportional to the indicated number of interactions or interaction strengths of ligand-receptor pairs. **(C)** Circle plots showing the inferred intercellular communication networks for WNT, and non-canonical WNT (ncWNT) signaling. The edge width is proportional to inferred interaction strengths considering ligand-receptor pairs. Black arrowheads show directionality of the inferred interactions. For (C), the color code for cell types is identical to (B).

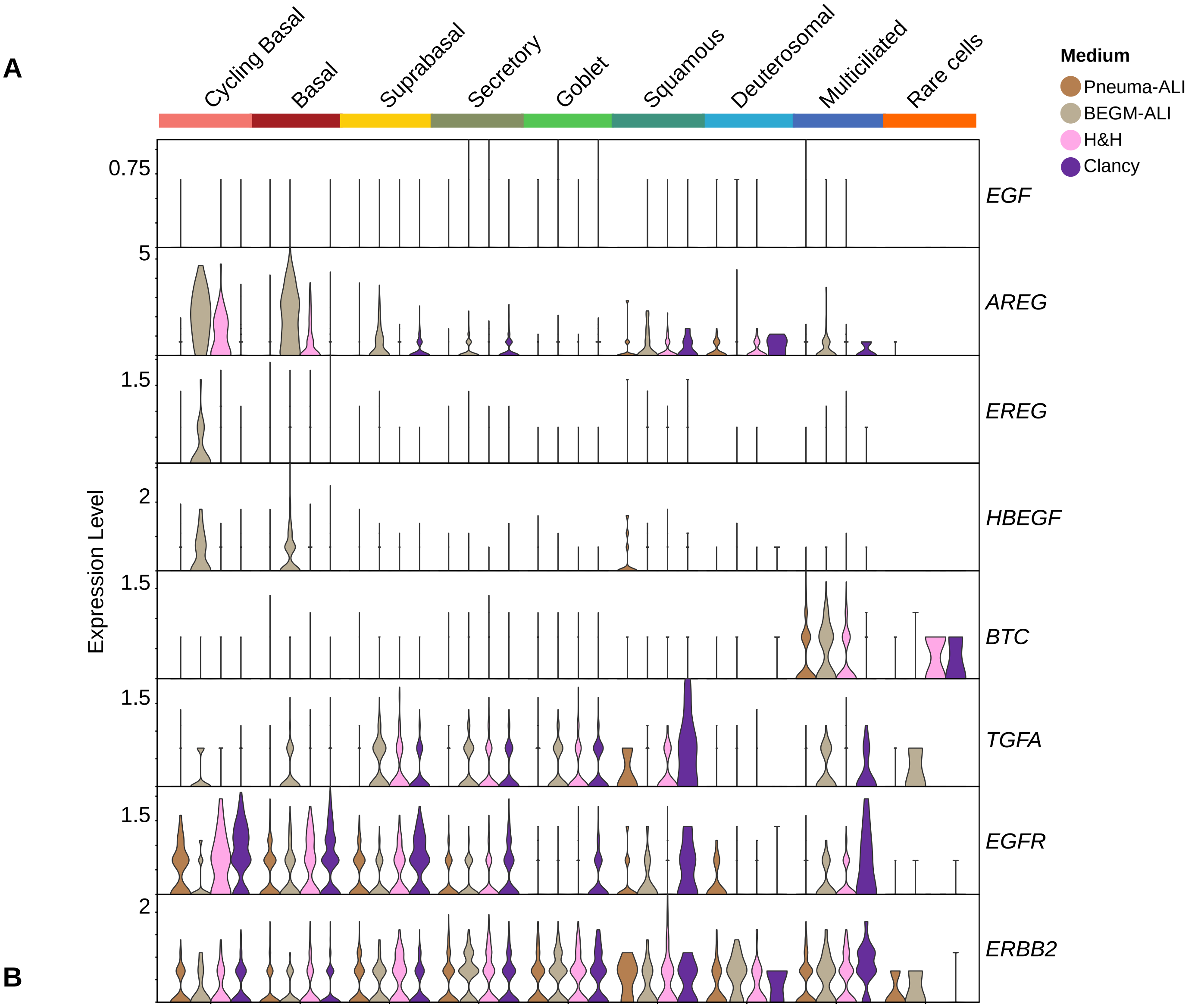


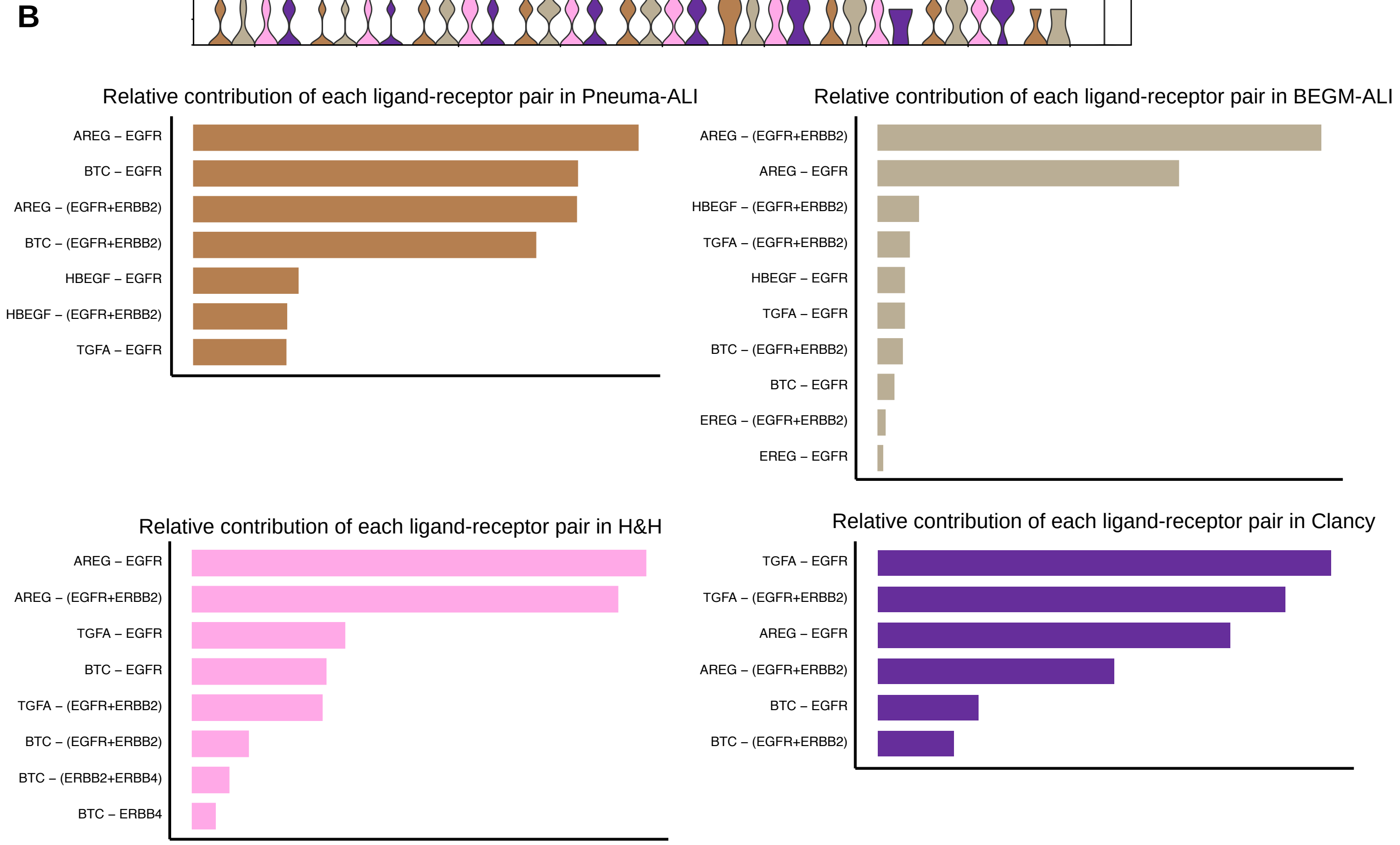


**Supplementary Figure 8:** EGF pathway member expression at ALI28 of bronchial epithelial cell differentiation.
**(A)** Violin Plots showing expression of ligands and receptors from the EGF pathway in 4 differentiation media. **(B)** Relative contribution of each Ligand-Receptor pair to interaction probabilities, in each differentiation medium. **(C)** Communication probabilities for each Ligand-Receptor pair, for each combination of sending-receiving cell type, in BEGM-ALI and Pneuma-ALI

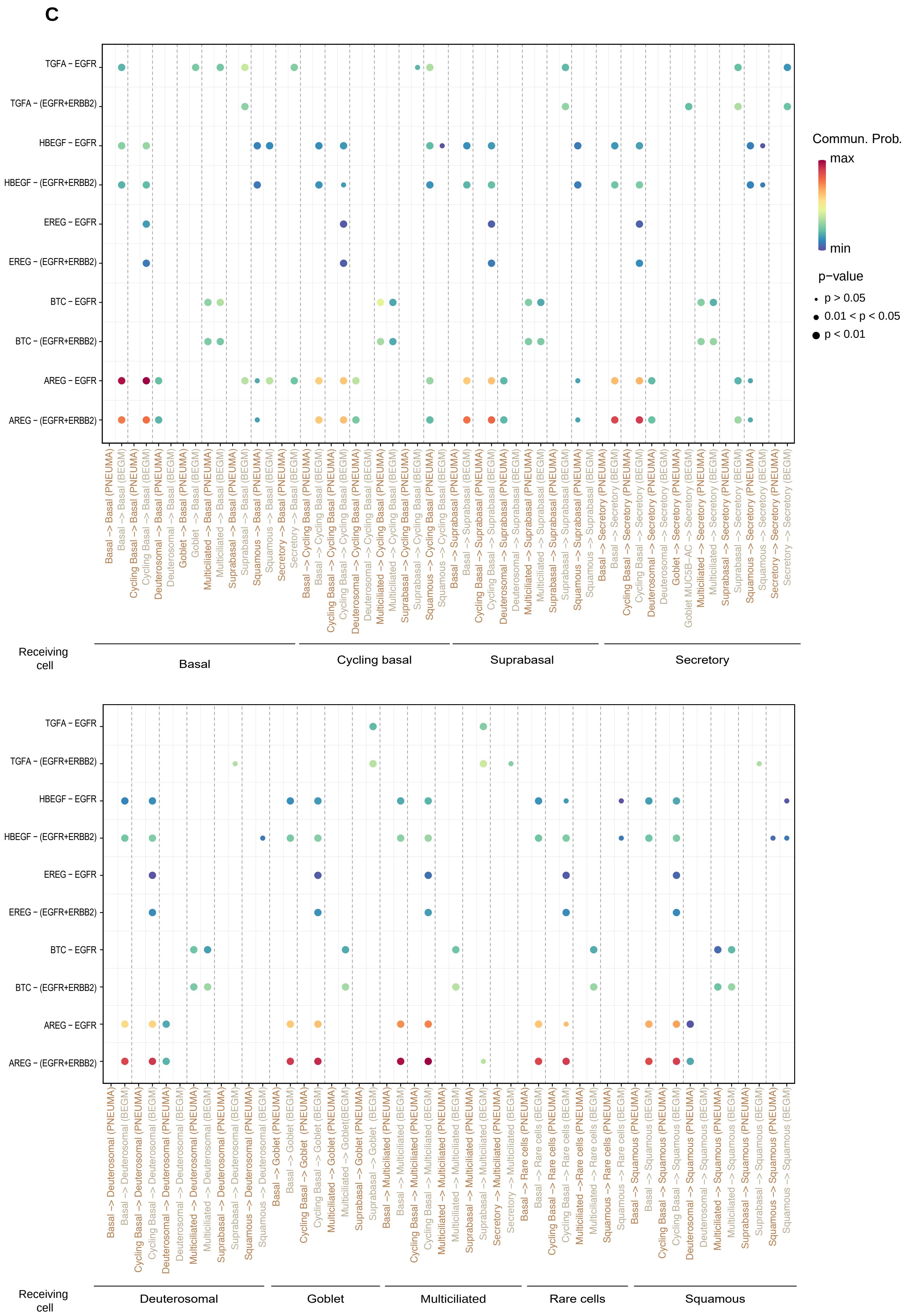


**Supplementary Figure 8-continued**

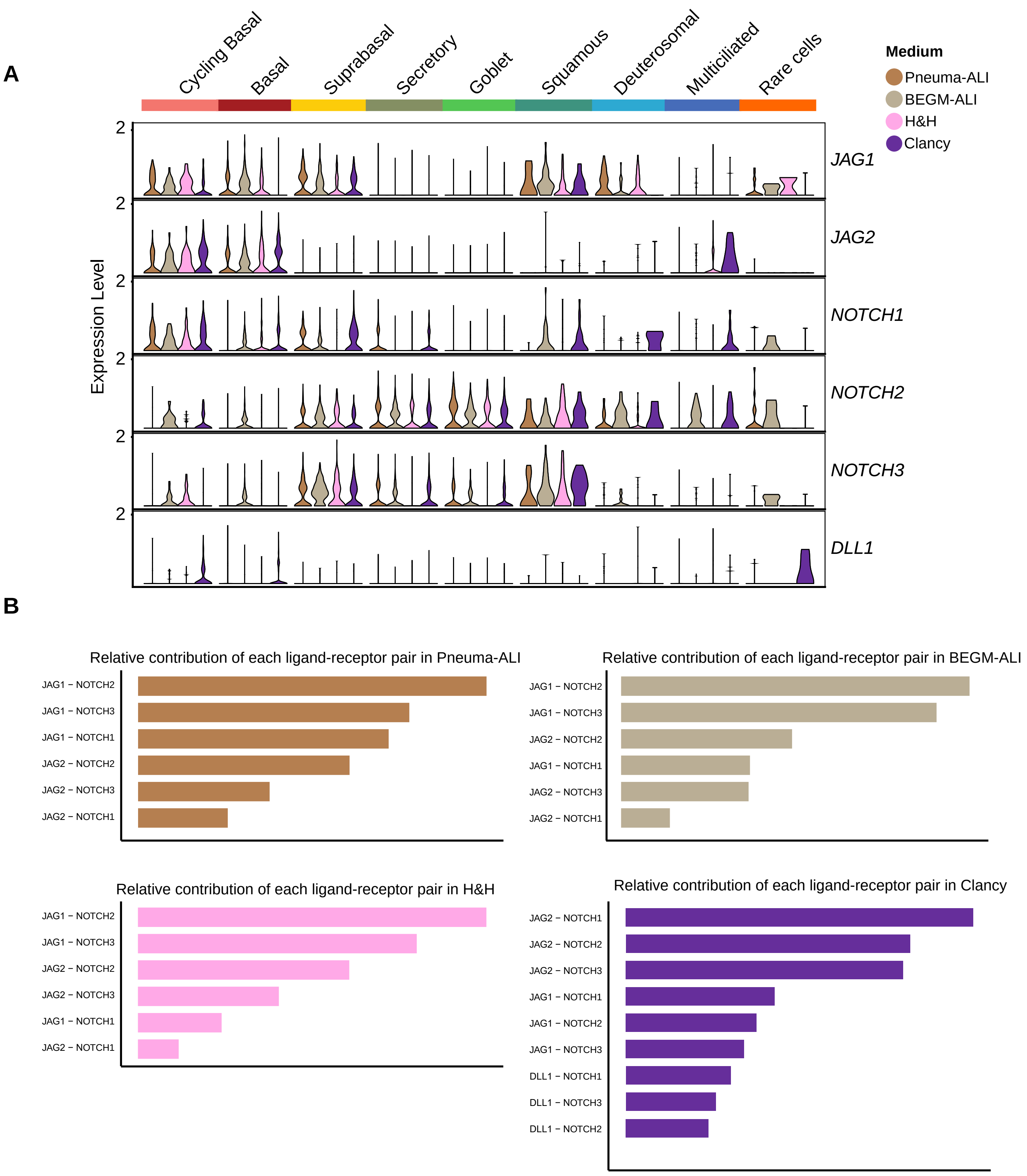


**Supplementary Figure 9: NOTCH pathway member expression at ALI28 of bronchial epithelial cell differentiation.** **(A)** Violin Plots showing expression of ligands and receptors from the NOTCH pathway in 4 differentiation media. **(B)** Relative contribution of each Ligand-Receptor pair to interaction probabilities, in each differentiation medium. **(C)** Communication probabilities for each Ligand-Receptor pair, for each combination of sending-receiving cell type, in BEGM-ALI and Pneuma-ALI.

C

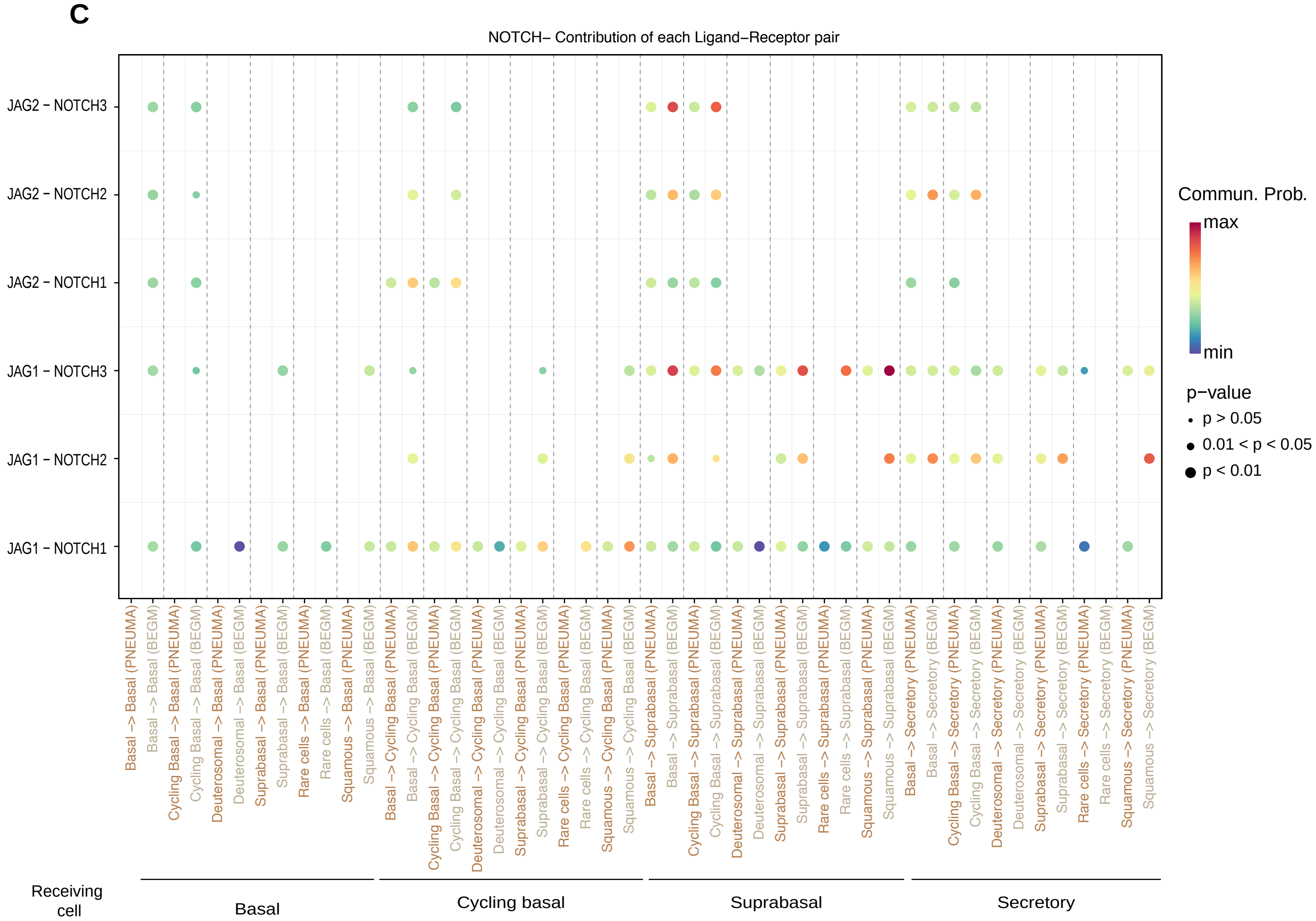


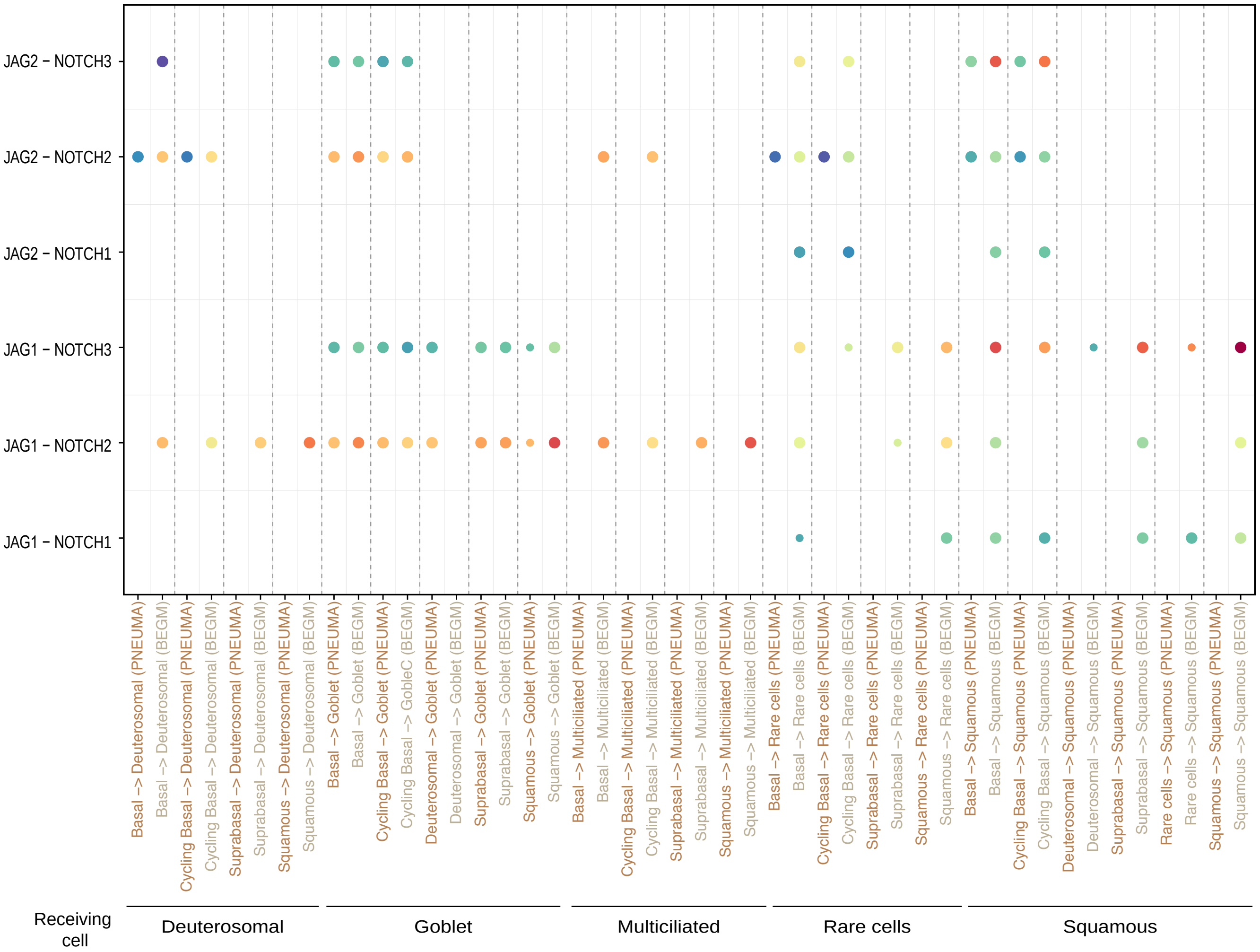


**Supplementary Figure 9-continued**

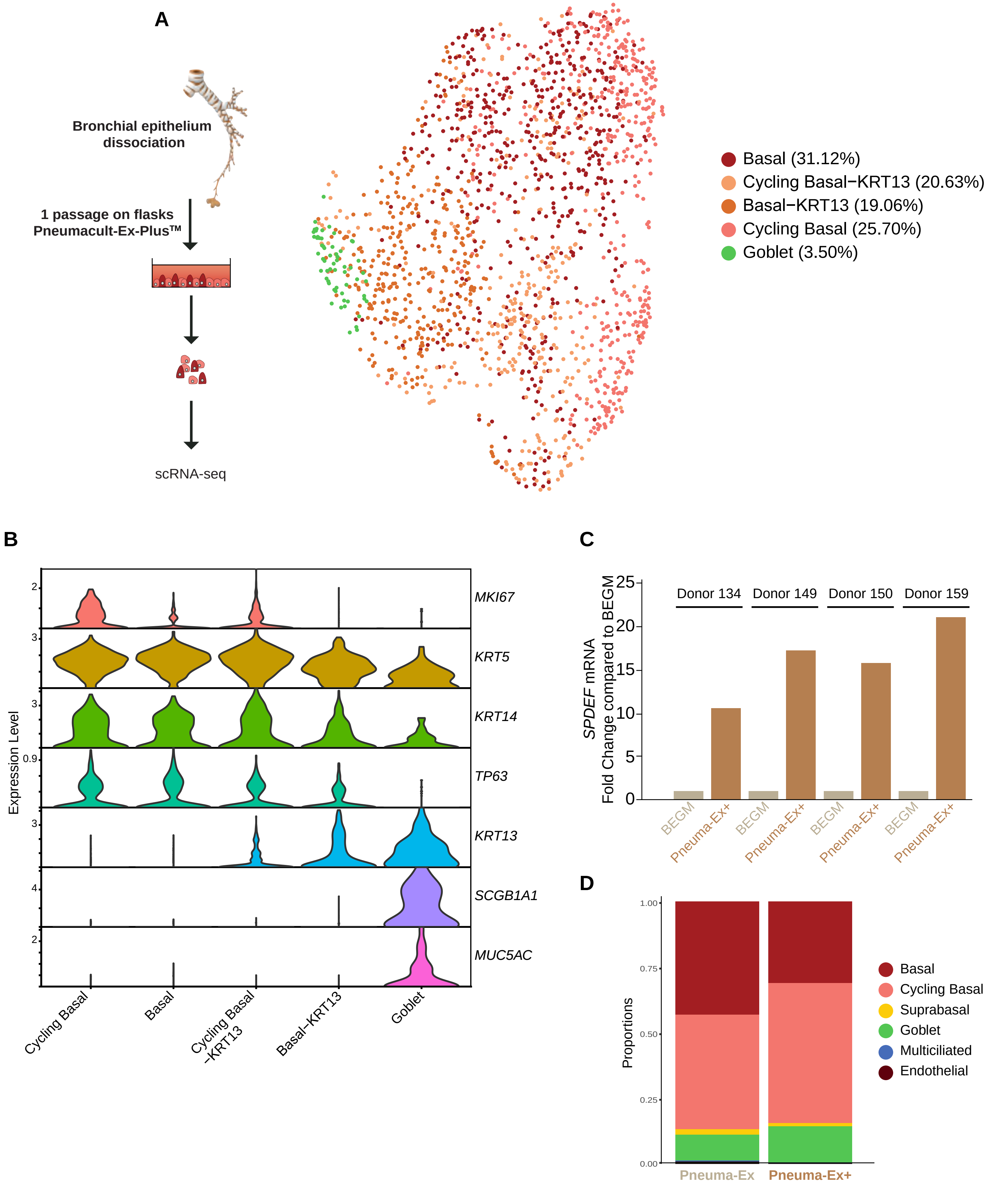


**Supplementary Figure 10: Distribution of cell types during the first cell propagation on plastic flasks.**
**(A)** UMAP representation of 1716 cells analyzed after dissociation from plastic flasks, at the time of TranswellTM seeding, from 2 independent HBEC cultures from 2 donors. **(B)** Expression level of a selected set of gene markers in the 5 detected cell clusters. **(C)** SPDEF expression (qPCR) for 4 independent HNEC cultures, during proliferation on plastic on flasks, after one passage in either BEGM or Pneuma-Ex+. **(D)** Quantification of cell type proportions from the single-cell data obtained from 3 independent HNEC cultures during proliferation on plastic on flasks, after one passage in either Pneuma-Ex or Pneuma-Ex+.

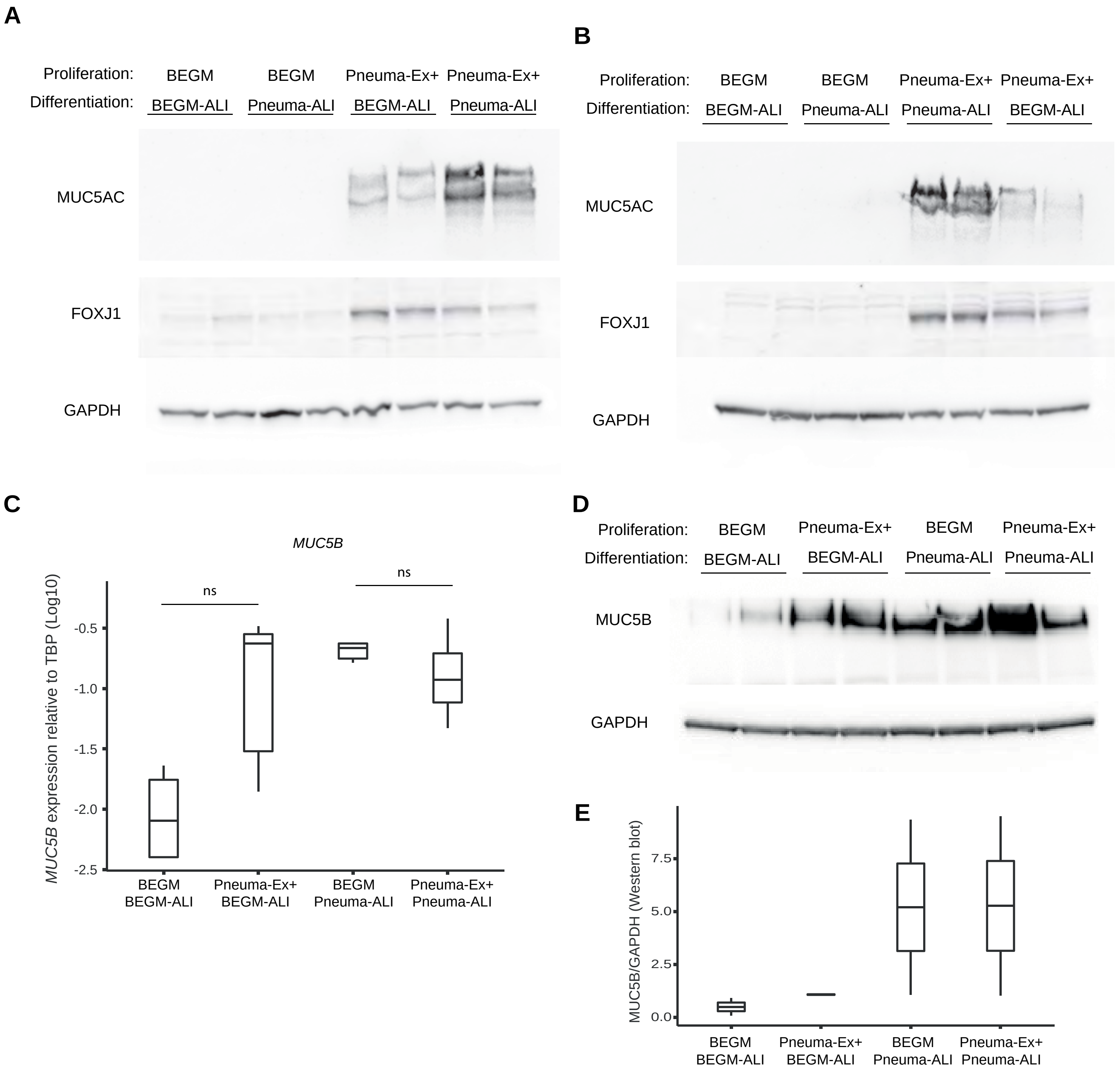


**Supplementary Figure 11: Effect of 2 proliferation media and 2 differentiation media on MUC5AC, FOXJ1 and MUC5B expression after differentiation of nasal epithelial cells.**

Western blot for MUC5AC and FOXJ1 on ALI cultures after full differentiation in the indicated media. GAPDH is used as loading control. **(A) and (B)** correspond to replicates from 2 independent cell cultures from 2 independent donors, which are distinct from the one presented in figure 4. For each condition, two lanes were loaded, each with an independent Transwell™ membranes from the same culture. **(C)** Expression of MUC5B (qPCR) on ALI cultures after full differentiation of nasal epithelial cells in the indicated media. Data shown are the values for log10(2^Ct(MUC5B-TBP)) for 3 independent cultures from 3 donors. p-values are the results of Mann-Whitney U test. **(D)** Western blot for MUC5B on ALI cultures after full differentiation in the indicated media. GAPDH is used as loading control. Protein extracts were from the same donor as in Figure 4. **(E)** Quantification of the MUC5AC signal from (C), p-values are the results of Welch's t-test.

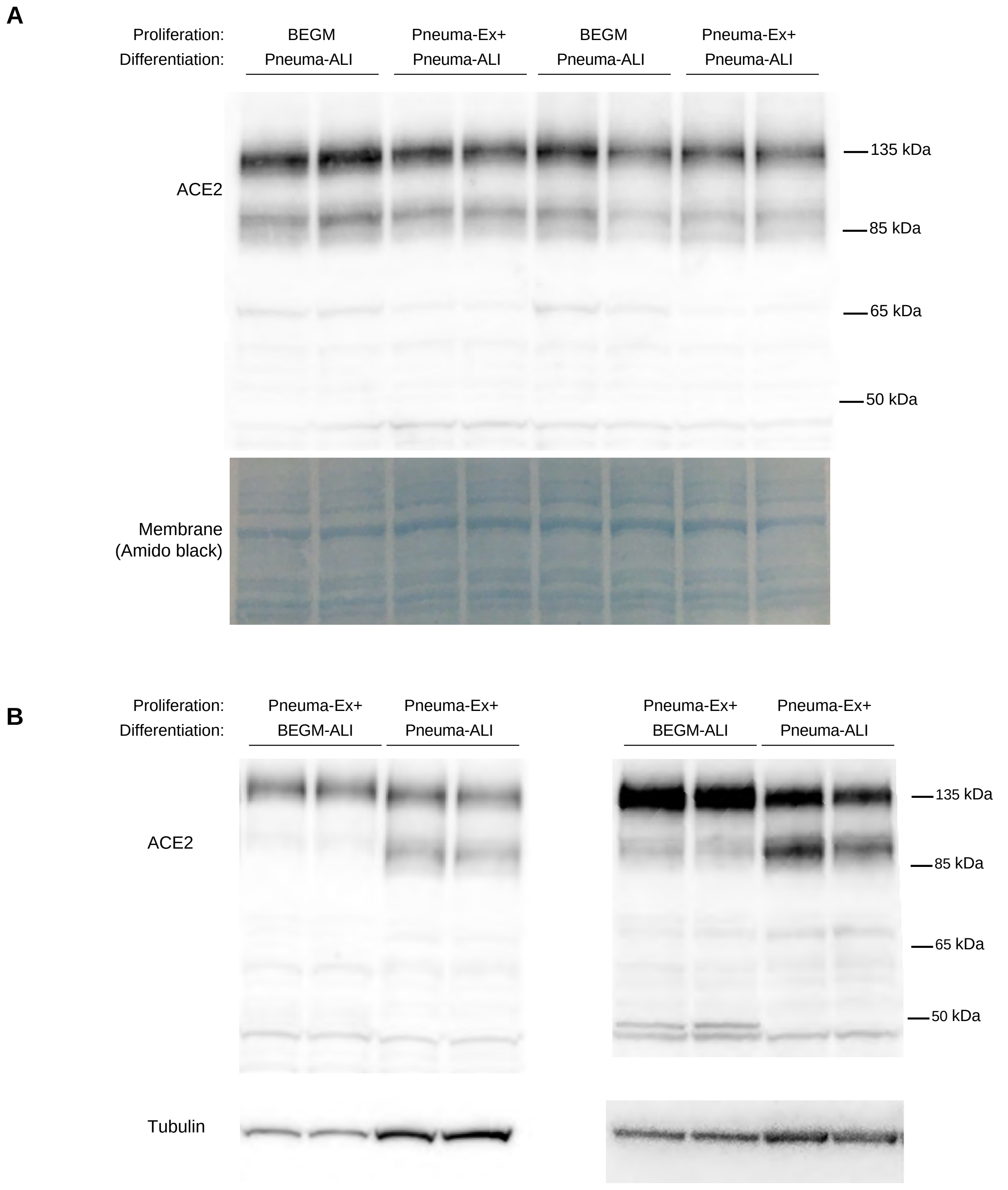


**Supplementary Figure 12: Effect of 2 proliferation media and 2 differentiation media on ACE2 expression profiles content after differentiation of nasal epithelial cells.**

**(A)** Western blot for ACE2 on HNEC ALI cultures after proliferation in either BEGM or Pneuma-Ex+, followed by full differentiation in Pneuma-ALI in the indicated media. Amido black staining of the membrane was used as loading control. Left and right parts of the blots correspond to replicates from 2 independent cell cultures from 2 independent donors, which are distinct from the ones presented in figure 7. For each condition, two lanes were loaded, each with an independent well from the same culture. **(B)** Western blot for ACE2 on HNEC ALI cultures after proliferation in Pneuma-Ex+, followed by full differentiation in either BEGM-ALI or Pneuma-ALI. Left and right panels correspond to replicates from 2 independent cell cultures from 2 independent donors, which are distinct from the ones presented in figure 7. For each condition, two lanes were loaded, each with an independent well from the same culture.

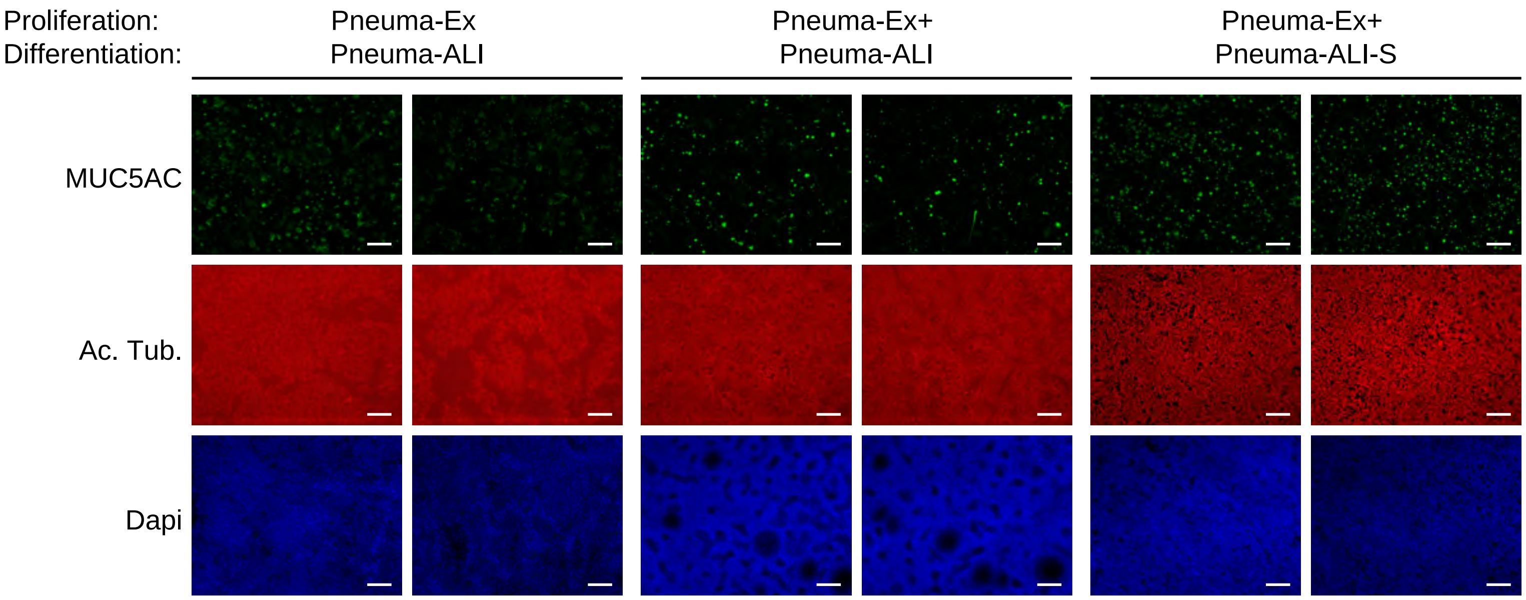


**Supplementary Figure 13: Effect of an alternative differentiation medium on epithelial composition after ALI differentiation of nasal epithelial cells.**

Additional images of immunostaining for MUC5AC and acetylated alpha-tubulin on HNEC ALI cultures after full differentiation in the indicated media. For each condition, two wells from the same culture batch are shown. Nuclei were stained with DAPI and shown in blue on the merge images; scale bars: 30 µm.
